# From Mutations to Disease: Computational Analysis and Interpretation of GPCR-Associated Pathogenicity

**DOI:** 10.1101/2024.06.17.599437

**Authors:** João Paulo L. Velloso, Alex G. C. de Sá, David B. Ascher

## Abstract

G protein-coupled receptors (GPCRs) perform critical roles in numerous physiological processes and their mutations are, therefore, associated with various human diseases. Hence, understanding the molecular consequences of pathogenic mutations in GPCRs is essential for elucidating disease mechanisms and developing effective therapeutic strategies. In this study, we employed computational approaches to explore the impact of mutations on GPCRs using two distinct datasets: ClinVar and MutHTP. We first evaluated the performance of available pathogenicity predictors. Beyond that, we used statistical analysis to identify key characteristics of mutations in GPCRs leading to diseases. We first evaluated available computational predictors, such as SIFT, PolyPhen-2, PROVEAN, ESM1b, and AlphaMissense in classifying GPCR mutations. During this task, we observed that all predictors performed with reliability when assessing GPCR mutations leading to diseases in the ClinVar dataset. On the other hand, when dealing with the MutHTP dataset, all predictors demonstrated poor performance, emphasising the importance of dataset characteristics and the need for comprehensive evaluation when selecting mutation predictive tools for GPCR analysis. The statistical analysis of mutations on GPCRs and disease development suggests that mutations occurring in conserved regions or regions with stronger intermolecular interactions are more likely to disrupt protein function and contribute to disease pathogenesis. Additionally, regarding our analysis, we also obtained insights into the importance of hydrophobic interactions and hydrogen bonding patterns in mutations in GPCRs and pathogenicity. Overall, our study enhances our understanding of the molecular mechanisms underlying GPCR-associated diseases and provides valuable insights for future research and clinical diagnostics in this field.

## 1 BACKGROUND

G protein-coupled receptors (GPCRs) are crucial for numerous vital physiological processes, including the regulation of cell division and proliferation, modulation of neuronal activity, control of ion transport across cell membranes and within organelles, maintenance of homeostasis, and the modulation and alteration of cell morphology (Salon et al., 2011). Since their discovery in 1948 (Ahlquist, 1948), GPCRs have been the subject of extensive research. Important findings include but are not limited to the ability of GPCRs to couple to multiple classes of G-proteins (Gudermann et al., 1996), the existence of a spectrum of conformational states between active and inactive conformations (Weis & Kobilka, 2018), receptor oligomerisation (Milligan et al., 2019), allosteric binding sites (Christopoulos et al., 2004), biased signalling (Wootten et al., 2018) and modulation by both orthosteric and allosteric endogenous ligands (Adan & Kas, 2003; van der Westhuizen et al., 2015).

These GPCR genome-wide traits are susceptible to deleterious modifications, such as mutations, exposure to chemicals, or abnormal signalling, which can lead to various acute or chronic human diseases (Salon et al., 2011). Examples of the impacts caused by these modifications include strokes associated with A2a-adenosine receptor mutations (Chen et al., 2007; Duan et al., 2009), asthma linked to mutations in the β2-adrenergic receptor (Kawakami et al., 2004), cardiovascular diseases related to mutations in the β1-adrenergic receptor (Drake et al., 2006), and depressive disorders, associated to monoaminergic GPCRs (Senese et al., 2018). In addition, the indispensable role of GPCRs in biology extends to possible involvement in the development of pathophysiological conditions, such as pulmonary edema during SARS-CoV-2 infections (Abdel Hameid et al., 2021).

Among various types of modifications, Single Nucleotide Variants (SNVs) – which can cause missense mutations – stand out as the most frequent in humans (Collins et al., 1997; Mooney, 2005). SNVs refer to single nucleotide mutations occurring within the DNA coding region, leading to alterations in the protein sequence. These alterations can result in changes to protein stability, structure, or function, as well as impact the strength of interactions within protein complexes and with drugs (Kulandaisamy et al., 2020). These SNVs can have profound functional implications (Wang et al., 2024) and have been statistically correlated to several disease phenotypes (Cline & Karchin, 2011). For instance, SNVs in GPCRs have been implicated in conditions, such as dominant and recessive obesity (involving melanocortin receptors), nephrogenic diabetes insipidus (associated with vasopressin receptor mutations), retinitis pigmentosa (involving rhodopsin mutations), familial exudative vitreoretinopathy (linked to frizzled receptor mutations), and female infertility (related to mutations in the follicle-stimulating hormone receptor) (Zalewska et al., 2014).

The most common alterations resulting from mutations in GPCRs are either loss of function (LOF) or gain of function (GOF). As a result, GPCR mutations are associated with a broad spectrum of diseases and variations in drug response, offering insights into critical structure-function relationships. Over 600 LOFs and nearly 100 GOFs have been characterised for GPCRs across 55 genes, associated with over 66 different human monogenic diseases (Schoneberg & Liebscher, 2021; Schoneberg et al., 2004).

LOF mutations in GPCRs typically involve the blockade of signalling even when the receptor interacts with its agonist(s) (Thompson et al., 2024). These missense mutations, associated with LOF, can result in insertions or deletions, frameshifts, or partial or complete gene deletions (Thompson et al., 2024). Approximately, 80% of LOF mutations alter protein folding, leading to the retention of misfolded receptors within cells, where they may aggregate and cause cell death (Park, 2019). Even if the mutant GPCR reaches the cell surface, it may still interfere with ligand binding or G-protein coupling (Schoneberg & Liebscher, 2021). LOF mutations are predominantly inherited, with misfolding often attributed to factors, such as impaired chaperone binding or receptor instability (Schoneberg et al., 2004). The GPCR misfolding might occur due to a series of factors, including the inability to bind essential chaperones or general receptor instability. One example of mutation capable of causing misfolding is the missense mutation (Cys271Arg) in the melanocortin-4 receptor. This mutation causes a disruption of interactions on the disulphide bridge between TM3 and the extracellular loop, resulting in receptor malfunction and is linked to obesity (Venkatakrishnan et al., 2013). Misfolding in the GPCRs can also result in proteins that retain function but that for reasons of misallocation alone cannot function normally (Ulloa-Aguirre & Michael Conn, 2011).

In contrast, GOF-associated mutations confer autonomous signalling to the receptor, even in the absence of a ligand. These mutations are mainly acquired. Germline transmission is rare because of a disease-limited lifetime or reproductive inability (Schoneberg et al., 2004). It is important to mention that constitutive activity in GPCRs was observed for more than 60 wild-type receptors for biogenic amines, nucleosides, lipids, amino acids, peptides, and proteins from different species, including humans and commonly used laboratory animal species. (Seifert & Wenzel-Seifert, 2002). However, any alteration in this basal activity can lead to diseases (Seifert & Wenzel-Seifert, 2002). The first mutation causing GOF in GPCRs to be discovered involved a class A GPCR, the gene encoding human RHO pigment of the retinal rods (Nathans & Hogness, 1984). This mutation causes retinitis pigmentosa (RP), a photoreceptor degeneration that affects 1.5 million people worldwide (Schoneberg et al., 2004). On the other side, related to class B, we can mention Jansens’s metaphyseal chondrodysplasia, which is caused by a GOF on the gene encoding the parathyroid hormone-parathyroid hormone-related peptide receptor. This disease is associated with severe hypercalcemia and hypophosphatemia (Schipani et al., 1995).

Various sequence- and structure-based computational tools were developed to enable the characterisation of the effects of SNVs on proteins – e.g. SIFT (Ng & Henikoff, 2001), PolyPhen-2 (Adzhubei et al., 2010), and PROVEAN (Choi & Chan, 2015). More recently, we also had some development of two deep learning models, ESM1b (Brandes et al., 2023), and AlphaMissense (Cheng et al., 2023), which were built for predicting the effects of missense variants in the human genome. Nevertheless, the study of missense mutations in membrane proteins has its own challenges. Membrane proteins are peculiar because of their physicochemical requirements. As a result, they have very strict particularities, which are associated with the fact that part of or the whole protein is embedded into a phospholipid bilayer (the cellular membrane). This fact usually leads to poor performance of universal predictive tools, as they are typically trained on the entire range of cellular proteins (Gromiha & Ou, 2014; Kulandaisamy et al., 2020). Because of that, we have some endeavours focused on developing tools focused on predicting the effect of mutations on membrane proteins (e.g., Pred-MutHTP (Kulandaisamy et al., 2020) and BorodaTM (Popov et al., 2019)).

Given the medical importance of GPCRs among membrane proteins, this study aims to explore a range of predictive tools for assessing the impact of mutations on GPCRs. We also performed structural- and sequence-based analysis, aiming to improve the comprehension of mutations in GPCRs (Figure 1). As already mentioned, understanding the effects of mutations in GPCRs is crucial due to their central role in mediating cellular signalling and their extensive involvement in various physiological processes. By analysing these predictive computational tools, we have gained valuable insights into the mechanisms underlying GPCR activation and how mutations can disrupt these processes. The findings from our study hold significant promise to enhance our exploration and comprehension of GPCRs. Moreover, the results derived from our analysis will serve as a foundation for future research in the field of clinical diagnostics for patients with GPCR-related conditions.

**Figure 1:**
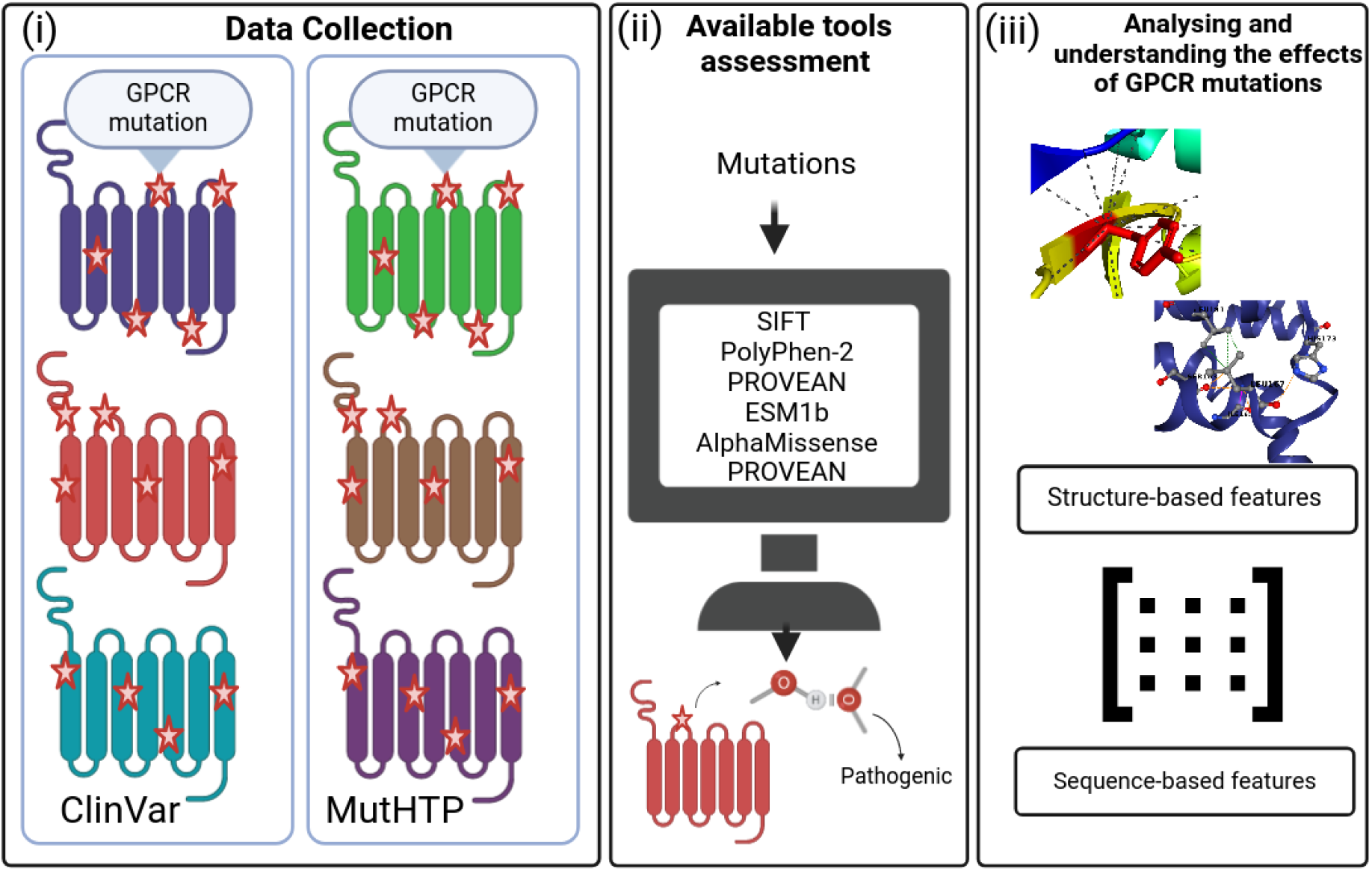
General workflow. i) Collection and curation of GPCR data on pathogenic and benign GPCR mutations. Two datasets were curated and analysed: ClinVar and MutHTP. (ii) Exploration of a range of predictive tools available for assessing the impact of mutations on GPCRs: SIFT (Ng & Henikoff, 2001), PolyPhen-2 (Adzhubei et al., 2010), PROVEAN (Choi & Chan, 2015), ESM1b (Brandes et al., 2023), and AlphaMissense (Cheng et al., 2023). iii) Modelling, statistically analysing and understanding the effects of mutations using various types of features, e.g., graph-based signatures, amino acid substitution scoring matrices (Blosum62, PAM30 and special amino acids), relative solvent accessibility, residue depth, AA index, molecular interactions.

## 2 RESULTS

### 2.1 Data collection

We compiled data from two sources: MutHTP (Kulandaisamy et al., 2018) and ClinVar (Landrum et al., 2014), each offering distinct perspectives on GPCR mutations and their links to diseases.

ClinVar comprehends 4,706,039 mutation entries, in which mutations are associated with their clinical significance (i.e., uncertain significance, benign, likely benign, pathogenic, likely pathogenic). After filtering it for only GPCR mutations, 57,042 were found, where 53,443 are missense mutations. Only 776 GPCR missense mutations in ClinVar have available experimental structures. We have considered two states of GPCR activation for these mutations – i.e., we have protein structural data comprehending all 776 mutations for the active state, and structural data for all 776 mutations for the inactive state. Across these mutations, 354 are labelled as disease-causing (i.e., pathogenic), whereas 422 are labelled as neutral (i.e., benign). These missense mutations are distributed across 216 GPCR proteins, where the five most frequent GPCRs include P08100 (n=78 mutations), P30518 (n=41), P32245 (n=29), P41180 (n=27), and P22888 (n=22). Most of the receptors belong to class A (n=609). Considering the other classes, 80, 65, and 22 mutations belong to the classes C, B, and F, respectively. In terms of regions, 38 mutations are located inside the loop region, 108 in helix 1, 88 in helix 2, 140 in helix 3, 86 in helix 4, 108 in helix 5, 122 in helix 6, and 86 in helix 7.

MutHTP, in turn, covers 206,390 transmembrane missense mutations, the mutations in this dataset are correlated to portions of the body (for example, upper aerodigestive tract, skin, ovary, or thyroid) and disease classes (Ex: cancers, nervous system diseases). After filtering out the GPCR mutations, we extracted 12,441 mutations. Among them, 12,014 are missense mutations, where 4,751 mutations have available experimental structures. 4,018 out of these mutations were classified as related to cancer. Like ClinVar, we have considered two states while dealing with GPCR structures – i.e., we have structural information of all 4,751 mutations for their respective active state, and, we have the structural information of all 4,751 mutations for their respective inactive state. Out of 4,751 GPCR mutations, 352 are labelled as “Neutral” (from now on, they will be defined as benign), and 4,399 are labelled as disease-causing (from now on, they will be defined as pathogenic). Across the pathogenic mutations, 4,111 are associated with cancer. These missense mutations are distributed across 370 GPCRs. The five most frequent GPCRs (and their number of mutations) include P30518 (n=92 mutations), O00222 (n=78), P24530 (n=75), Q6W5P4 (n=70), and P08100 (n=62). Considering GPCR classes, most of them belong to class A (n=3,909). In addition, the number of mutations for the classes C, B, and F, are 376, 367, and 99, respectively. Considering the different regions of the receptor, 320 mutations happen inside loops, 608 in helix 1, 572 in helix 2, 726 in helix 3, 526 in helix 4, 782 in helix 5, 715 in helix 6, and 502 in helix 7.

### 2.2 Computational assessment of mutation impact on GPCRs

In this section, we explore a range of predictive tools to assess the impact of mutations on GPCRs, retrieved from two distinct sources: ClinVar and MutHTP. Whereas ClinVar integrates mutation data from multiple sources, including peer-reviewed literature, clinical testing laboratories, locus-specific databases, and expert-curated databases, MutTHP mutations are mostly correlated to cancer. For each data set, we applied two different analyses, which are described next.

First, we assessed the performance of existing computational methodologies in predicting the pathogenicity of mutations on GPCRs. The considered tools for this step were SIFT (Ng & Henikoff, 2001), PolyPhen-2 (Adzhubei et al., 2010), PROVEAN (Choi & Chan, 2015), ESM1b (Brandes et al., 2023), and AlphaMissense (Cheng et al., 2023). These tools were selected in our evaluation because they are the most widely used and are well-established computational methods for predicting the pathogenicity of mutations (Dong et al., 2015). We have also considered ESM1b and AlphaMissense because they are two recent models based on deep learning. ESM1b and AlphaMissense are known to achieve great predictive performance as they were trained on a massive amount of data (Brandes et al., 2023; Cheng et al., 2023). Nevertheless, it is important to mention neither of these tools is specialised in GPCRs, possibly lacking GPCR-related characteristics and particularities while modelling them for machine (and deep) learning pipelines.

In this part of the assessment, various metrics – including the Matthews correlation coefficient (MCC), accuracy, area under the curve score (AUC), weighted F1 score, sensitivity and specificity (see Supplementary Material for more information about these metrics) – are considered in our analysis to comprehensively and complementarily estimate the performance of each prediction tool. By systematically applying these predictive tools to our dataset of GPCR mutations, the aim is to evaluate their predictive performance and identify the most reliable tools for assessing the impact of mutations on GPCRs.

Second, we applied predictive computational methods to comprehensively understand the underlying correlation between pathogenesis and mutations in GPCRs. This analysis is supported by statistical tests to identify molecular drivers of diseases upon mutations in GPCRs. In the Methodology section, we describe how we performed this statistical analysis of the effects of mutations on GPCRs using different descriptive features. This integrated approach allowed us to identify key changes caused by mutations in GPCRs that cause GPCR-related disorders and provided a foundation for further exploration in this field.

#### 2.2.1 ClinVar data

##### 2.2.1.1 Benchmark, analysis and understanding of predictive mutagenesis tools on GPCR mutations

The evaluation of computational tools that are commonly used to predict the pathogenicity of mutations for the ClinVar dataset offered useful insights into their performance. As is shown in Table 1 and Figure 2, AlphaMissense was established as the best tool for accurately predicting the impact of mutations on GPCRs in the ClinVar dataset, consistently presenting superior performance metrics including accuracy, MCC, F1-score, AUC, sensitivity and specificity. Additionally, ESM1b, SIFT, and PROVEAN have also demonstrated good performance, further adding to the toolkit of accurate mutation predictive tools in the evaluated task. PolyPhen-2 scored slightly lower performance across all evaluated metrics, but even then, it can be considered a reliable tool for the tested scenario. All evaluation metrics for classification are detailed in the Supplementary Materials.

**Figure 2:**
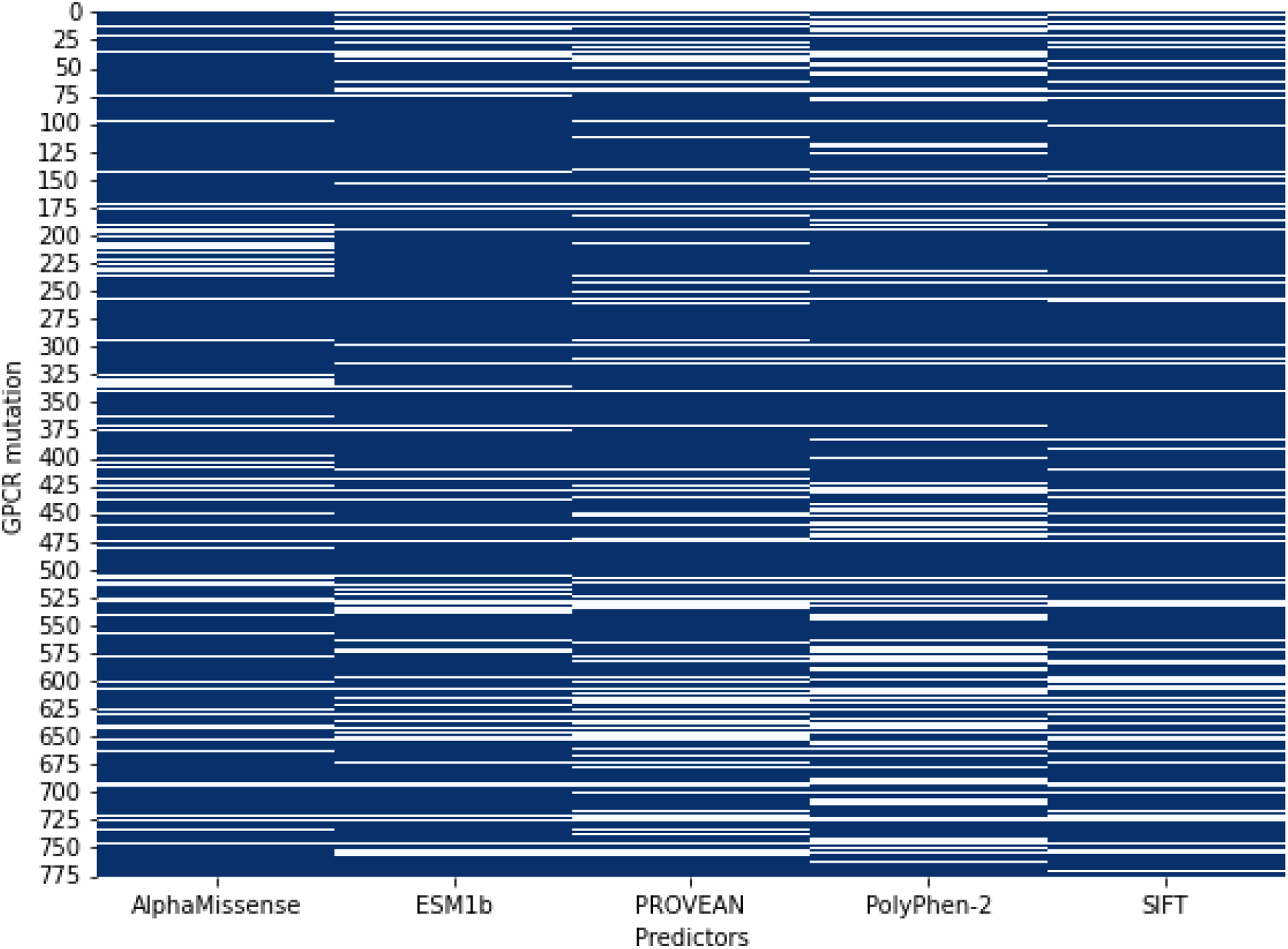
Heatmap depicting the accuracy of various predictors in identifying the clinical significance of variants. Each row in the heatmap represents a distinct GPCR mutation, while each column corresponds to a specific predictor used in the analysis. Blue cells indicate instances where the predictor correctly identified the clinical significance of the variant for that GPCR mutation, while white cells indicate the respective predictor’s classification errors. This visual representation allows for a comprehensive comparison of the performance of different predictors across multiple GPCR mutations.

**Table 1:** Predictive performance of predictive tools for assessing the impact of GPCR mutations in ClinVar data.

| Method | Accuracy | MCC | F1-score weighted | AUC | Sensitivity | Specificity |
| --- | --- | --- | --- | --- | --- | --- |
| **AlphaMissense** | 0.82 | 0.68 | 0.86 | 0.85 | 0.90 | 0.89 |
| **SIFT** | 0.78 | 0.61 | 0.78 | 0.79 | 0.96 | 0.64 |
| **PolyPhen-2** | 0.72 | 0.50 | 0.72 | 0.73 | 0.90 | 0.57 |
| **ESM1b** (used the threshold of -7.5, lower than -7.5 means the mutation is pathogenic) (https://doi.org/10.1038/s41588-023-01465-0) | 0.81 | 0.65 | 0.81 | 0.82 | 0.90 | 0.74 |
| **PROVEAN** (considered a score higher than -2.5 to be benign and lower than or equal to -2.5 to be pathogenic). | 0.76 | 0.55 | 0.75 | 0.77 | 0.92 | 0.62 |

In Figure 2, we present a heatmap depicting the accuracy of various tested predictors in identifying the clinical significance of variants. It provides a visual comparison of the predictor accuracy across different GPCR mutations. Each row in the heatmap represents a distinct GPCR mutation (classification), while each column corresponds to a specific predictor used in the analysis. Blue cells indicate instances where the predictor correctly identified the clinical significance of the variant for that mutation (benign or pathogenic). White cells indicate the predictor’s misclassifications. As observed by our benchmark it is possible to check that AlphaMissense has the higher number of correct predictions, presenting an accuracy of 0.82. ESM1b presented a slightly lower accuracy of 0.81. The remainder of the tools were also performed with the reliability of this data set, accuracies of 0.78, 0.72, and 0.76 were observed for SIFT, PolyPhen-2, and PROVEAN, respectively.

In addition to evaluating the overall performance of our predictors, we conducted a focused analysis on cases where the AlphaMissense predictor made incorrect predictions. This analysis aimed to determine whether other predictors could compensate for AlphaMissense errors. To visualise this, we generated a second heatmap in Figure S1 that includes only the GPCR mutations where AlphaMissense failed to correctly identify the clinical significance of the variant. In other words, where AlphaMissense outputs wrong predictions across pathogenic and benign labels. Figure S1 follows the same format as our initial analysis, with each row representing a distinct GPCR mutation and each column representing a tool other than AlphaMissense. A blue cell indicates a correct prediction by the corresponding predictor for that GPCR mutation. This focused visualisation helps to identify which alternative predictors are most reliable in compensating for the errors made by AlphaMissense. Furthermore, we were able to gain insights into their relative strengths and weaknesses, by pointing out visually which tools predicted properly when AlphaMissense failed. In summary, our benchmark analysis can guide the development of more robust predictive models by highlighting predictors that consistently provide accurate classifications when others fail. Considering this, it is possible to observe that both SIFT and PolyPhen-2 represented good alternatives to be combined with AlphaMissense. They displayed overall good predictive performance in cases where AlphaMissense failed.

To confirm this trend, we generated confusion matrices to provide a detailed view of how each predictor performs in terms of classifying the clinical significance of variants that AlphaMissense failed to predict properly. These matrices can be seen in Figure 3. Each subplot within Figure 3 represents a confusion matrix for a specific predictor, considering the cases where the AlphaMissense made an incorrect prediction. By examining these matrices, we can see the distribution of correct and incorrect predictions for each predictor. For instance, a higher count of true benign (top-left cell) and true pathogenic (bottom-right cell) cells indicates better performance, while higher counts in the false pathogenic (top-right cell) and false benign (bottom-left cell) cells highlight areas where the predictor struggles. Considering these premises, we can verify that PolyPhen-2 and SIFT (Figure 3, subplots C and D, respectively) are indeed the best tools to be combined with AlphaMissense, because they have a higher count of true benign and true pathogenic. SIFT has a count of 11 and 60 for true benign and true pathogenic, respectively (Figure 3D). PolyPhen-2, in turn, accounted for 17 and 55 of true benign and true pathogenic, respectively. We suggest that the user could combine the SIFT and PolyPhen-2 predictions to better identify pathogenic missense GPCR mutations with AlphaMissense, assigning different weights to them. For example, provided with the predictive results, we recommend AlphaMissense predictions having a higher weight (e.g., 3 or 5), and the SIFT and PolyPhen-2 with a lower weight (e.g., 1 or 2 each). If we consider a majority voting with the assigned weights, predictive results might be more trustworthy.

**Figure 3:**
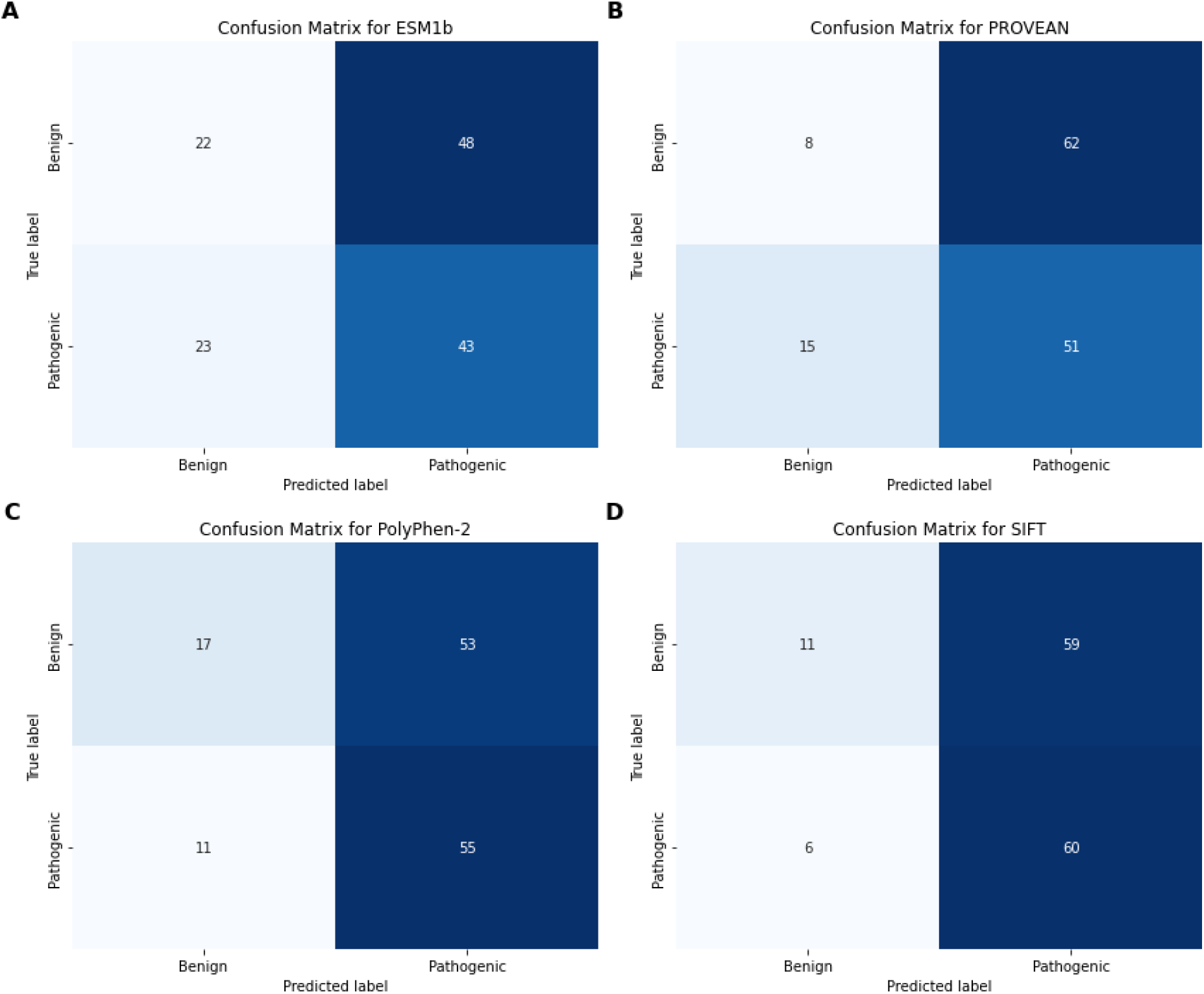
Confusion matrices for the A: ESM1b, B: PROVEAN, C: PolyPhen-2, and D: SIFT predictors, showing their performance in classifying the clinical significance of variants. Each subplot presents a matrix with true labels on the y-axis and predicted labels on the x-axis. The cell values indicate the number of GPCR mutations in each category. These matrices illustrate the accuracy of each predictor, with a focus on cases where AlphaMissense made incorrect predictions. True Benign (Top-left cell): GPCR mutations correctly predicted as Benign. True Pathogenic (Bottom-right cell): GPCR mutations correctly predicted as Pathogenic. False pathogenic (Top-right cell): GPCR mutations incorrectly predicted as pathogenic. False Benign (Bottom-left cell): GPCR mutations incorrectly predicted as benign.

Furthermore, we analysed the performance of each predictor considering the following information about the mutations presented in the ClinVar dataset: UniProt ID, GPCR classes, phenotype, and cellular position. For this analysis histograms were generated, which provide a clear and straightforward representation of the predictors’ performance for each evaluated case (see Figures S2, S3, S4, and S5). Each histogram subplot in Figures S2, S3, S4, and S5 corresponds to one predictor and shows the accuracy for each class (see Supplementary materials for more details about this metric). The x-axis of each histogram represents the different information used for grouping the mutations per predictive tool, while the y-axis indicates the predictive tools’ accuracy. By examining these histograms, we can identify trends and patterns in the performance of each predictor, e.g.: consistency across the grouping of characteristics of the mutation (predictors that show a relatively uniform distribution of correct predictions across all classes are likely more consistent and reliable); and characteristic-specific performance (variability in the height of the bars across different characteristic for a given predictor can indicate that the predictor performs better for some grouping than others). These results are crucial for understanding the strengths and weaknesses of each predictor in specific and different contexts.

For instance, when analysing the performance of predictors by grouping the dataset using UniProt IDs in Figure S2, it becomes evident that AlphaMissense exhibits significant variability in its performance. For example, AlphaMissense correctly predicts 61% of cases for the UniProt ID P30968 (Gonadotropin-releasing hormone receptor), whereas it achieves a 95% accuracy for P30518 (Vasopressin V2 receptor). In contrast, the other tools demonstrate lower variability in their prediction success rates compared to AlphaMissense. Notably, for the GPCR UniProt ID O75899 (Gamma-aminobutyric acid type B receptor subunit 2), there is a marked drop in performance across all tools when compared to the average performance for all groups. This suggests that certain predictors may be more sensitive to specific protein contexts, highlighting the importance of evaluating predictive tools in a nuanced, context-dependent manner. Considering the variability observed for AlphaMissense, the other tools could support more reliable predictions since the variability on them is much lower.

When analysing the accuracy by grouping the data according to GPCR classes A, B, C, and F (Figure S3), we observe that the performance is notably higher for GPCR class F (Frizzled/Smoothened) across all tools. Surprisingly, despite most available data on GPCRs being related to Class A, the performance for this class is not as high as expected. This trend highlights that the predictive tools are not specifically tailored for GPCRs, indicating a potential gap in their design and optimisation. This lack of specialisation suggests that these tools may not fully capture the unique structural and functional nuances of GPCRs, particularly for the more extensively studied Class A. Consequently, there is a pressing need for the development and refinement of predictive models that are specifically optimised for the diverse classes of GPCRs. Such tailored predictive tools could significantly enhance the accuracy and reliability of predictions, ultimately improving our understanding and ability to target these critical receptors in clinical and therapeutic contexts.

We also analysed the accuracy of the tools by grouping the mutations according to their disease phenotype as shown in Figure S4. Among the various analyses conducted on the ClinVar dataset, this one showed the highest variability between groups. For instance, when considering the disease Epileptic Encephalopathy, which is characterised by severe, early-onset seizures and significant neurodevelopmental impairments (Hwang & Kwon, 2015), all the tools displayed low accuracy, with PolyPhen-2 achieving the lowest accuracy ratio (i.e., at 16%). In significant contrast, for some phenotypes, the tools achieved accuracies as high as 100%, for example, ESM1b, PROVEAN, and SIFT accuracies reached 100% for diabetes insipidus-related mutations. PolyPhen-2 accuracy reached 100% for mutations related to Hypogonadotropic hypogonadism, Body mass index quantitative trait locus 20, and autosomal dominant hypocalcemia 1. Besides, SIFT accuracy also reached 100% for mutations related to Body mass index quantitative trait locus 20, and Glucocorticoid deficiency 1. This trend reflects the challenge of predicting the clinical significance of mutations associated with complex phenotypes and suggests that the tools may not be equally effective across different disease contexts. The substantial variability in performance highlights the necessity for phenotype-specific calibration of these predictive models. Enhancing the accuracy of these tools for diverse phenotypes is crucial, as it would lead to more reliable interpretations of genetic mutations, ultimately improving patient diagnosis and personalised treatment strategies.

At last, we analysed the tools’ accuracy according to cellular localisation (non-cytoplasmic domain, cytoplasmic domain, and transmembrane) (Figure S5). AlphaMissense and ESM1b show high performance across all localisations, achieving up to 85% accuracy in transmembrane regions. Notably, AlphaMissense, as the best method, consistently scores well, reaching 85% accuracy in non-cytoplasmic domains and 82% in transmembrane regions. PROVEAN and PolyPhen-2, while slightly lower compared to AlphaMissense, and ESM1b still demonstrate robust performances. These results highlight the overall high predictive accuracy of AlphaMissense and ESM1b, especially in transmembrane regions. The trend reiterates the importance of cellular localisation in prediction accuracy and suggests that while predictors perform well across different localisations, possible enhancements are still needed for all domains. This chart emphasises the necessity of refining prediction models for specific cellular contexts to achieve consistent high accuracy across all domains.

Overall, our analysis of ClinVar data highlights significant variability in the accuracy when considering distinct categorisations of the mutations. Understanding these variations is crucial for improving the accuracy and reliability of predictive models in clinical genomics, as it allows for the identification of predictors that may require further refinement or alternative approaches for specific protein groups or cases.

##### 2.2.1.2 Statistical analysis and interpretation of GPCR mutations

Moving forward, we aimed to identify features (i.e., molecular drivers) capable of distinguishing between pathogenic and benign GPCR mutations within the ClinVar dataset. Features exhibiting a stronger correlation with the outcome were subjected to detailed exploration, providing insights into the underlying molecular mechanisms that lead to disease development. As outlined in the Methodology section, our approach involved statistical analysis based on a diverse array of features to uncover molecular drivers of mutations associated with GPCR-related disorders. To achieve this, we assessed the predictive usefulness of these features in mutation classification, using machine learning modelling (see the Methodology section). Additionally, we took advantage of Shapley Additive Explanations (SHAP) (Lundberg & Lee, 2017) to identify the most influential features for classifying GPCR mutations into pathogenic or benign. SHAP plots highlighting the most important molecular drivers for predicting GPCR mutations are depicted in Figure 4 (also Table S1), supporting the interpretation of GPCR pathogenicity predictions.

**Figure 4:**
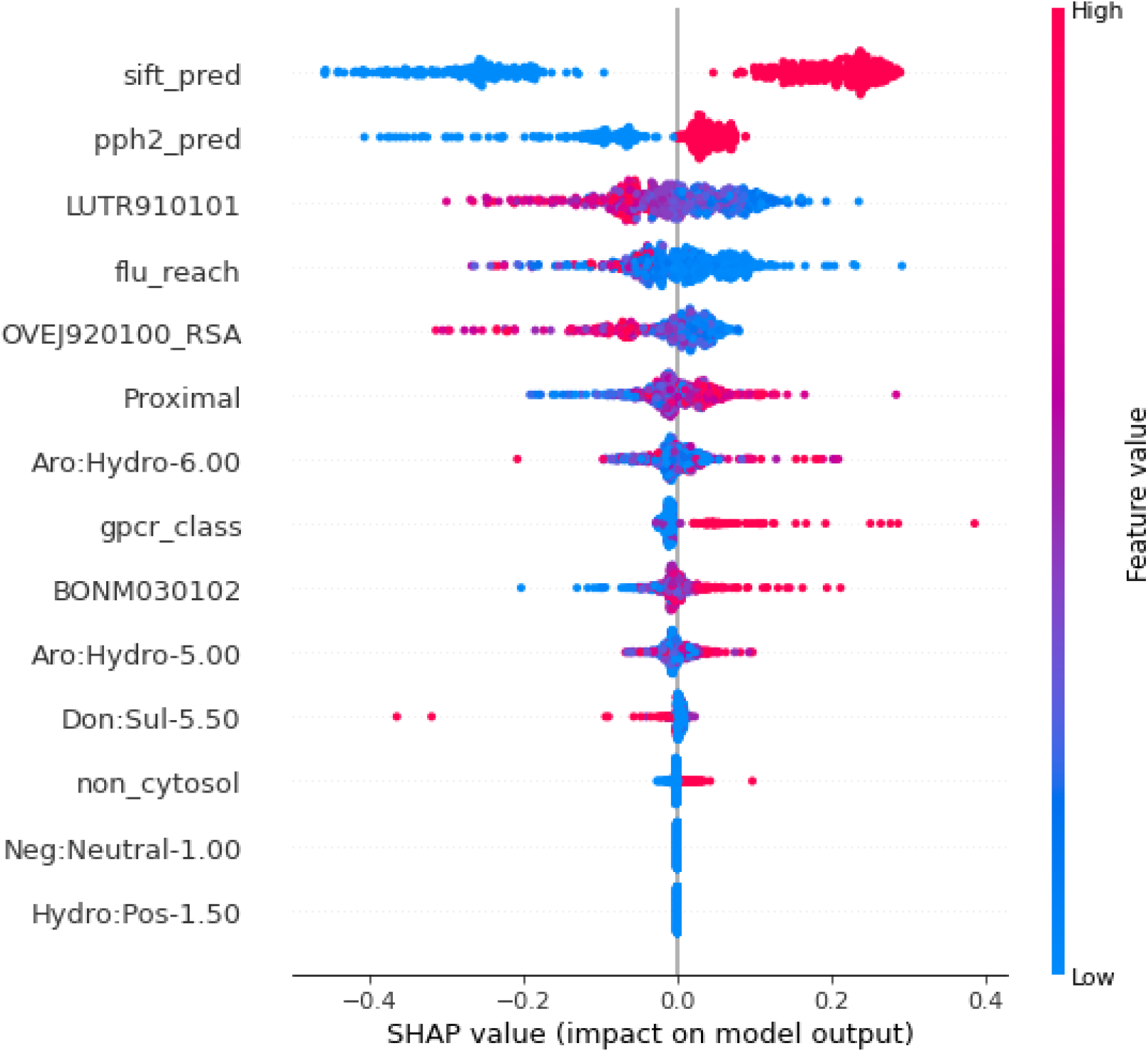
SHAP feature importance plot for ClinVar dataset. The features that represent higher performance in distinguishing between pathogenic and benign mutations are represented in descending order of importance on the vertical axis. The points represent the training dataset. High values are represented in red colour, while points with low values are presented in blue. Points to the left influence the prediction to be benign (negative SHAP values), and points, to the right, are pathogenic (positive SHAP values). Different input features affect the output of the respective classification.

The two most important features are those based on already published predictors of the possible impact of amino acid substitution on proteins, *sif_pred* (i.e., SIFT predictions) and *pph2_pred* (i.e., PolyPhen-2 predictions), see Figure 4. As mentioned before, both predictors could properly classify the mutations into benign or pathogenic for the ClinVar dataset, reassuring their overall importance in our assessed meta-classifier.

After *sif_pred* and *pph2_pred*, our analysis discovered three important features extracted from AAindex (Kawashima et al., 2008): LUTR910101 (Luthy et al., 1991), OVEJ920100_RSA, and BONM030102 (Figure 4). OVEJ920100_RSA (AAINDEX: OVEJ920105, (Overington et al., 1992)) refers to an environment-specific amino acid substitution matrix for inaccessible residue. It indicated the substitution probability (lower values, lower probability) for inaccessible residues (buried). According to Overington et al. (1992), generally for all residues a buried position is more conserved than a surface position. This supports the results found in the SHAP plot, where low values of OVEJ920100_RSA are correlated to pathogenic mutations, which is also corroborated by the violin plot (see Figure S6A). In summary, these results map those mutations happening in regions with a lower probability of changes of amino acid (i.e., more conserved regions) tend to lead to disease.

The LUTR910101 index, in turn, refers to a structure-based comparison table (Luthy et al., 1991). Specifically, it indicates the possibilities of changes of amino acids in regions outside of secondary structure classes that are neither Alpha helix nor beta-strand. Low values are related to more conserved residues for the mentioned case, and positive values are related to less conserved residues. According to the SHAP plot, for this feature, low values are correlated to mutations being classified as pathogenic. An observation that is also corroborated by the violin plot (see Figure S6B). These results indicate that if a mutation occurs in a conserved residue, it tends to lead to disease. This observation points out the significance of evolutionary conservation in protein function and highlights the risk of deleterious effects of mutations disrupting conserved regions. Such mutations are more likely to interfere with essential protein-protein interactions, structural stability, or functional domains, ultimately contributing to disease manifestation (Xiong et al., 2022).

BONM030102 (Boniecki et al., 2003) was the last AAindex identified by SHAP. It is a feature characterising neighbour atom contact properties in biomolecular interactions. Particularly, it refers to the Quasichemical Statistical Potential (QSP) for the intermediate orientation of interacting side groups, representing the interaction energy between two side groups, such as amino acid residues in proteins (Boniecki et al., 2003). QSP is calculated based on statistical considerations of their relative orientations, considering the distribution of possible side group orientations and the probabilities of observing different configurations. Higher values of this feature for the intermediate orientation of interacting side groups typically indicate stronger interactions or more favourable configurations between the biomolecular entities under consideration (Boniecki et al., 2003). According to the SHAP plot in Figure 4, higher values are correlated with pathogenicity, which is also confirmed by the violin plot in Figure S6C. This observation aligns with previous findings suggesting that mutations occurring in regions involved in stronger intermolecular interactions or more favourable configurations are more likely to disrupt protein function and contribute to disease pathogenesis (Livesey & Marsh, 2022).

Three descriptors coming from graph-based signatures (see Supplementary Materials for more details) were shown to be important for the task of identifying pathogenic or benign GPCR (Figure 4): Aro:Hydro-6.00 (mapping pairs between aromatic atoms and hydrophobic atoms at the distance of 6 angstroms), Aro:Hydro-5.00 (mapping pairs between aromatic atoms and hydrophobic atoms at the distance of 5 angstroms), Don:Sul-5.50 (mapping pairs between hydrogen bond donor atoms and Sulphur group atoms at the distance of 5.50 angstroms). As mentioned before, these features represent the quantification of pairs of pharmacophoric regions within a defined distance threshold surrounding the mutation site. The features Aro:Hydro-6.00 and Aro:Hydro-5.00 are very similar (Figure S7A, and S7B), having just a difference in the cut-off distance to measure the number of Aromatic-Hydrophobic pairs, 6 and 5 angstroms respectively. These two features showcase the importance of hydrophobic surfaces, including the ones from aromatic rings (de Araujo et al., 2022) to keep the proper folding and function of membrane receptors in general. Hydrophobic interactions contribute to an important role in membrane insertion and folding of GPCRs (https://doi.org/10.1016/S1043-9471(05)80049-7). Any changes in the hydrophobic core can cause the protein to lose function. The feature Don:Sul-5.50 (Figure S7C) represents a more complex case, and it represented a feature of less impact for the classification. This feature importance could be correlated to the importance of hydrogen bond interactions for protein function (Bertalan et al., 2020; Venkatakrishnan et al., 2019). Overall, the identification of these graph-based signatures highlights the significance of hydrophobic interactions and hydrogen bonding patterns in GPCR stability and function, and the importance of observing them when classifying mutations on GPCRs.

#### 2.2.2 MutHTP data

A similar methodology to ClinVar data was applied to MutHTP data to derive its analysis. Contrastingly, different patterns and trends were observed in MutHTP analysis, including the best predictors found for this dataset, as indicated in the Supplementary Results.

According to the Supplementary Results for MutHTP data, predicting, understanding and explaining the effects of GPCR mutations leading to cancer is complex and therefore challenging to be computationally identified (and interpreted) by machine learning pipelines and models. We assume that this is caused by the various factors involving cancer development, such as the involvement of complex biological pathways, the influence of the tumour microenvironment, and the need to distinguish mutations that are really involved and mutations that are present but have no impact on cancer development. Given these challenges, the development of specialised tools is of major importance. As seen through these results, new tools should consider information regarding the mutations, such as subcellular localisation, class of the GPCR, the specific GPCR involved in the disease, and disease phenotypes caused by the mutation. These predictive tools may ultimately contribute to more effective diagnostics and personalised treatment strategies for cancer patients, especially if their results are combined or redesigned to consider GPCR’s inherent characteristics.

#### 2.2.3 Performance dichotomy on GPCR-mutation pathogenicity prediction between ClinVar and MutHTP

The contrasting results observed between the two datasets highlight the importance of dataset characteristics and the performance of predictive tools. This suggests that while AlphaMissense may excel in certain aspects, however, caution should be exercised in relying solely on one prediction tool, and further improvements may be needed to enhance the reliability of GPCR mutation classification. This also suggests the importance of integrating multiple predictive tools to enhance prediction reliability.

One of the possible reasons for this dichotomy is that the MutHTP data is mostly composed of mutations leading to cancer. As mentioned before, predicting cancer is particularly challenging due to the involvement in cancer of complex biological pathways, the dynamic and heterogeneous nature of the tumour microenvironment, and the difficulty in distinguishing mutations that have an impact on the cancer development and mutations that are present, but do not have an impact in the disease onset.

Furthermore, when comparing both datasets, we found 314 entries that were common to both. Out of these common entries, 224 had consistent classifications between the two datasets (90 classifications were divergent). The disagreement between the two datasets could be attributed to various factors. A possible factor is the difference from the data sources utilised for compiling mutation information, which may result in variations in the mutations documented between the datasets. Additionally, discrepancies in curation methods, such as criteria for inclusion or exclusion of mutations, and the interpretation of literature findings, might have led to divergent annotations. Variances in annotation standards, scope, and focus of the datasets, as well as updates and revisions over time, further contribute to inconsistencies in mutation-disease mappings. Moreover, human error may introduce an extra level of discrepancies. Overall, these factors highlight the complexities and challenges associated with GPCR mutation annotation for phenotype understanding and, also, the importance of cautious interpretation when utilising mutation data from different sources. The discrepancies help to explain the differences in predictive performance when testing available tools for the classification of mutations into pathogenic or benign classes. Further validation and comparison with additional data are essential to support these findings and ensure their applicability across diverse datasets. This further highlights the importance of improving predictors for GPCR analysis, emphasising how GPCRs represent a highly distinct family of proteins with unique functional characteristics.

## 3 CONCLUSIONS

In this study, we investigated the performance of predictive tools capable of differentiating mutations as pathogenic or benign when dealing with GPCR mutations. Additionally, we explored potential molecular mechanisms that can lead to disease upon mutations in GPCRs, employing statistical analysis. Two distinct datasets, MutHTP and ClinVar, were analysed, providing unique perspectives on GPCR mutations and their associations with diseases. The comprehensive analysis of these datasets highlights the complexities involved in predicting the impact of GPCR mutations and underscores the necessity for advanced computational tools in this domain. This study contributes valuable insights into the performance of existing tools and the potential for developing more precise methods for GPCR mutation analysis.

When we evaluated the various tools using the ClinVar dataset, which is characterised by its diverse collection of genetic mutations and their clinically annotated significance, encompassing a wide range of diseases, AlphaMissense emerged as the top-performing tool, demonstrating consistently high accuracy, MCC, F1-score, and AUC values, suggesting its efficacy in assessing mutation pathogenicity in this dataset. In contrast, the MutHTP dataset, primarily comprising mutations correlated with cancer, presented significant challenges in classification due to its imbalanced nature and complex mutation landscape.

Despite efforts to discern pathogenic from benign mutations using various predictive tools, including SIFT, PolyPhen-2, and AlphaMissense, the results exhibited limited reliability across different datasets, highlighting the challenging nature of GPCR mutation classification. The varying performance of predictive tools between the two datasets emphasises the importance of dataset characteristics and the need for comprehensive evaluation when selecting mutation predictive tools for GPCR analysis.

Going further we conduct analysis to check combinations of tools to support a more reliable analysis of GPCR mutations. Considering the ClinVar data, AlphaMissense, the top performer can be combined to SIFT and PolyPhen-2. These two tools manage to predict properly some cases where AlphaMissense failed. Following the same idea, considering the MutHTP data set, PROVEAN and SIFT could boost performance of the top performer, PolyPhen-2.

We also analysed the performance of each predictor considering the following information about the mutations presented in the ClinVar dataset: UniProt ID, GPCR classes, phenotype, and cellular position. Throughout these analyses, it was evident that there is a significant variability in the accuracy when considering distinct categorisations of the mutations. Understanding these variations is crucial for improving the accuracy and reliability of predictive models in clinical genomics, as it allows for the identification of predictors that may require further refinement or alternative approaches for specific protein groups. For example, surprisingly, despite most available data on GPCRs being related to Class A, the performance for this class is not as high as expected, when we considered both ClinVar and MutHTP data. We also observed that the accuracy varies greatly according to the receptor being analysed. It can be as high as 100% for the receptor UniProt ID P30968, and as low as 47% for the receptor UniProt ID O75899, both cases considering PolyPhen-2 performance on the ClinVar data set. Besides, the phenotype caused by the mutation can also lead to different accuracies. When considering the MutHTP for all predictors, the accuracies related to cancer mutations were the lowest when compared to other phenotypes. In this case, considering all tools, the lowest accuracy was 38% obtained for AlphaMissense and the highest was 69% for PolyPhen-2 when grouping the mutations by disease phenotype.

During our statistics analysis, trying to understand possible features driving the development of disease upon mutations, we could identify features that had higher impacts on distinguishing mutations on GPCRs as pathogenic or benign on the ClinVar data set. On this group of features, the ones that stood out were extracted from AAindex (LUTR910101, OVEJ920100_RSA, and BONM030102). Low values of LUTR910101 and OVEJ920100_RSA, indicative of conserved residues, were correlated with pathogenic mutations, highlighting the deleterious effects of mutations disrupting conserved regions and essential protein-protein interactions. Similarly, the Quasichemical Statistical Potential represented by BONM030102 underlined the possible correlation between the development of disease and mutations that occur in protein regions with a higher number of intermolecular interactions. Furthermore, we identified some graph-based signatures, such as Aro:Hydro-6.00, Aro:Hydro-5.00, and Don:Sul-5.50 that displayed importance for distinguishing mutations on GPCRs into pathogenic and benign. These signatures highlighted the critical role of hydrophobic interactions and hydrogen bonding patterns in GPCR stability and function, offering promising predictive features for classifying mutation pathogenicity. These findings provide valuable insights into the mechanisms underlying GPCR-associated diseases and illustrate the importance of computational approaches in elucidating mutation impact and guiding future research efforts aimed at therapeutic interventions. While performing the same analysis for the MutHTP dataset, the results were not very clear, and all features exhibited a notably low screening performance. The three most important features were DOSZ010101 (extracted from AAindex, 17998252, using an Amino acid similarity matrix based on the sausage force field, 11524370), Hbond (number of hydrogen bonds on the mutation site), and the graph-based signature Acc:Hydro-3.00 (this feature indicates the count of pairs of Hydrogen bond acceptors atoms and hydrophobic atoms within a maximum distance of 3 angstroms from the mutation site). These features’ importance is likely correlated to their association with hydrogen bonds and hydrophobic interaction formation, which are critical in stabilising protein structure and function (Bertalan et al., 2020; de Araujo et al., 2022; Venkatakrishnan et al., 2019). However, they did represent the same reliability as the features found for ClinVar data, given the challenges found in MutHTP (cancer-associated) data.

Furthermore, the discrepancies observed between the MutHTP and ClinVar datasets point to the challenges associated with mutation annotation and classification. Factors such as data sources, curation methods, and annotation standards contribute to inconsistencies, necessitating cautious interpretation when utilising mutation data from different sources. The substantial disagreement between the datasets illustrates the complexities inherent in GPCR mutation analysis in terms of disease phenotype identification.

Finally, due to the critical role of GPCRs in a wide range of physiological processes and their involvement in numerous diseases, including cancer, and considering the low performance measured of available tools when these tools were employed to the MutHTP dataset, improving predictors for GPCR is imperative. Accurate prediction of the effects of mutations on GPCRs may lead to a better understanding of disease mechanisms and the development of good, targeted therapies. By combining advanced computational tools and integrating multiple prediction approaches, researchers can gain deeper insights into the biological consequences of GPCR mutations and their association with disease. Additionally, further validation and comparison with additional datasets are crucial to verify findings and ensure the applicability of predictive models across diverse mutation landscapes. Specifically, incorporating features such as localisation, class, and protein type could significantly improve prediction accuracy. This study underscores the multifaceted nature of GPCR mutation analysis and the ongoing efforts to advance predictive capabilities in this field. Continued advancements in this area will enable more accurate disease predictions and potentially uncover novel therapeutic targets, ultimately contributing to a better understanding and treatment of GPCR-related disorders.

## 4 METHODOLOGY

The general workflow of this study is shown in Figure 1. The steps followed were: (i) Data collection and curation, which refers to collecting experimental data about GPCR mutations and disease onset in humans. Two datasets were analysed, ClinVar and MutHTP. (ii) Exploration of a range of predictive tools available for assessing the impact of mutations on GPCRs: SIFT (Ng & Henikoff, 2001), PolyPhen-2 (Adzhubei et al., 2010), PROVEAN (Choi & Chan, 2015), ESM1b (Brandes et al., 2023), and AlphaMissense (Cheng et al., 2023). iii) Modelling, statistically analysing, explaining and understanding the effects of mutations through various types of descriptive features – e.g. graph-based signatures, amino acid substitution scoring matrices (Blosum62, PAM30 and special amino acids), relative solvent accessibility, residue depth, AA index, molecular interactions – were performed with statistical methods. This third step in the proposed methodology is also important so that we can characterise the physical and biochemical changes caused by the evaluated mutations.

### 4.1 Data collection and curation

For this study, data were collected from two distinct sources: ClinVar (Landrum et al., 2014) and MutHTP (Kulandaisamy et al., 2018), each providing unique insights into GPCR mutations and their association with diseases. ClinVar was chosen for its comprehensive coverage, encompassing a diverse range of mutations, including both cancer and non-cancer variants, thereby providing a broader context for our analysis. Meanwhile, MutHTP was selected for its specialisation in mutations correlated with cancer when considering GPCRs, offering a focused dataset tailored for cancer evaluation.

It is worth noting that when considering MutHTP, every clinical significance was selected. On the other hand, to consider data coming from ClinVar, the following clinical significance filters were selected to create our dataset: “Likely benign”, “Benign”, “Benign/Likely benign”, “Pathogenic”, “Likely pathogenic”, and “Pathogenic/Likely pathogenic”. The mutations described as “Likely benign”, “Benign”, and “Benign/Likely benign, were labelled in our analysis as benign. The mutations described as “Likely pathogenic”, and “Pathogenic/Likely pathogenic” were labelled as pathogenic. The data retrieval process involved querying each database using an extensive list of UniProt IDs retrieved from GPCRdb (Isberg et al., 2016) covering all human GPCRs catalogued up to date. In addition, both datasets were subjected to thorough data extraction, ensuring the retrieval of relevant mutation information, including mutation type, gene/protein affected, and associated cancer types.

One of the aims of this study is to analyse the structural impact of these mutations on GPCRs. Because of that, only mutations covered on available GPCR structures were selected. The generation of features necessitated wild-type-like structures of the receptors present in the dataset. To achieve this, we employed inactive and active state GPCR structure-based models constructed using AlphaFold-Multistate. These GPCR structure models are conveniently accessible through GPCRdb (Pandy-Szekeres et al., 2023). It is important to emphasise that all generated models were based on the wild-type sequence. This step holds critical importance, especially considering that many GPCR structures available in the Protein Data Bank (Berman et al., 2000) contain mutations essential for increased stability and enabling experimental structure elucidation. By accessing models without these mutations on GPCRdb, we ensured consistency in our dataset. Additionally, we deliberately selected models without loops to mitigate the likelihood of bias resulting from regions with low AlphaFold pLDDT scores (Jumper et al., 2021). This careful selection process aimed to enhance the reliability and robustness of our evaluation of the impact of mutations on GPCRs.

The datasets with all the information and the predictions are in the supplementary material.

### 4.2 Exploration of a range of predictive tools available for assessing the impact of mutations on GPCRs

In this step, two separate analyses were conducted using the two mentioned datasets: one utilising the MutHTP dataset (more mutations correlated to cancer) and the other employing the ClinVar dataset (more diverse disease onsets). By conducting separate analyses using different datasets, we aim to comprehensively evaluate the predictive capabilities of these tools and assess their suitability for predicting the impact of mutations on GPCRs in different scenarios. The predictive performances of the following tools are studied: SIFT (Ng & Henikoff, 2001), PolyPhen-2 (Adzhubei et al., 2010), PROVEAN (Choi & Chan, 2015), ESM1b (Brandes et al., 2023), and AlphaMissense (Cheng et al., 2023). These tools were chosen for their ability to classify mutations as deleterious or benign, providing valuable insights into eventual disease onset. They are the state of the art regarding mutation pathogenicity analysis. The tool SIFT (Sorting Intolerant From Tolerant) uses sequence homology to predict the impact of single amino acid substitutions on protein function. SIFT was installed locally, version 6.2.1, NCBI BLAST 2.4.0+, database UniRef90. PolyPhen-2 (Polymorphism Phenotyping v2) combines information from protein sequence, evolutionary conservation, and structural features to classify substitutions as benign or damaging. PolyPhen-2 predictions were obtained through their web server, parameters: Classifier model “HumDiv”, Genome assembly “GRCh37/hg19”, Transcripts “Canonical”, Annotations “Missense”. PROVEAN assesses the impact of variations on protein function by comparing the sequence similarity between the query protein and its homologue. PROVEAN was installed locally, version 1.1.5, NCBI BLAST 2.4.0+, NCBI non-redundant protein database version 4. ESM1b uses a machine learning approach to classify variants as pathogenic or benign based on various features, including protein sequence, structure, and functional annotations. AlphaMissense utilises an ensemble of machine learning models trained on features derived from protein sequence, structure, and evolutionary conservation to classify mutations as deleterious or tolerated. ESM1b and AlphaMissense predictions were those available and precalculated for all human proteins. The statistics evaluation of this step was measured using MCC, accuracy, weighted F1 score, AUC, recall (sensitivity) and specificity. Predictions made by AlphaMissense categorised as ambiguous were excluded when calculating sensitivity and specificity. All evaluation metrics for classification are also detailed in the Supplementary Materials.

### 4.3 Modelling and statistical analysis of the effects of mutations on GPCRs

In our study, we undertook the task of comprehensively understanding the impact of mutations on GPCRs and their likely association with disease onset. To achieve this, we made a statistical analysis based on a comprehensive set of descriptors (or features) accounting for various protein properties, including descriptors of conservation and function, local residue environment, stability and dynamics, and protein interaction changes. By including these diverse aspects of protein structure and function, our aim is to identify potential molecular drivers of mutations in GPCRs leading to diseases. The analysis of these features provided a wholesome understanding of the complex interplay between mutations and GPCRs, shedding light on their likely implications for disease onset. General descriptions of these features are outlined in Table ***2*** (see Table S1 and Table S2 for more information about them).

**Table 2:** Sequence- and structure-based features derived for mutation analysis in both ClinVar and MutHTP datasets.

| Type | Name |
| --- | --- |
| Sequence-based | Amino acid substitution scoring matrices (Blosom62, PAM30 and special amino acids) |
|  | Relative solvent accessibility |
|  | Residue depth |
|  | AA index |
|  | Potential function energy calculations |
| Structure-based | Arpeggio - Molecular interactions |
|  | Graph-based signatures |
|  | Structural pattern mining approaches |
|  | Normal mode analysis |

Due to the highly imbalanced nature of the MutHTP dataset, which contains 4,851 GPCR mutations with only 355 classified as benign, we approached the data by separating it into benign and pathogenic sets. We then divided the pathogenic set into four equal, randomly selected subsets. Each subset of pathogenic mutations was combined with the benign set, creating four distinct sets of mutations. Statistical analyses of the effects of these mutations were conducted on all four sets, with the reported results reflecting the highest performance based on MCC values. We aimed to assess the correlation and predictive ability of certain features in classifying mutations. To achieve this, we employed three ML algorithms within scikit-learn V.1.2.1 (Pedregosa et al., 2011): Random Forest (Tin Kam, 1995), eXtreme Gradient Boosting (XGBoost) (Chen & Guestrin, 2016), and Extremely randomized trees (Geurts et al., 2006). All features were statistically analysed to evaluate their performance in classifying mutations in the classes pathogenic and benign. The statistics evaluation of this step was measured using MCC, accuracy, and weighted F1 score, AUC. All evaluation metrics for classification are also detailed in the Supplementary Materials. Additionally, we calculated SHAP values (SHapley Additive exPlanations) (Lundberg & Lee, 2017), and from these values, we generated plots. Which were used to point out the features that exhibit higher discriminative power. SHAP assigns an importance value to each feature with respect to the classification, offering a detailed understanding of the factors influencing the categorisation. It enables a better understanding of the classification of pathogenic and benign mutations. Within the SHAP summary plot, the visualisation represents the relationship between feature values and their impact on classifications. It illustrates how both low and high values of features are linked to either class (i.e., pathogenic or benign), aiding in understanding the predictor outputs. This visual representation enhances interpretability, facilitating the comprehension of the dynamics between input features and the model’s decision-making process.

## Supporting information

ClinVar variant summary

MutHTP variant summary

## REFERENCES

Abdel Hameid, R., Cormet-Boyaka, E., Kuebler, W. M., Uddin, M., & Berdiev, B. K. (2021). SARS-CoV-2 may hijack GPCR signaling pathways to dysregulate lung ion and fluid transport. Am J Physiol Lung Cell Mol Physiol, 320(3), L430–L435. 10.1152/ajplung.00499.2020

Adan, R. A., & Kas, M. J. (2003). Inverse agonism gains weight. Trends Pharmacol Sci, 24(6), 315–321. 10.1016/S0165-6147(03)00130-5

Adzhubei, I. A., Schmidt, S., Peshkin, L., Ramensky, V. E., Gerasimova, A., Bork, P., Kondrashov, A. S., & Sunyaev, S. R. (2010). A method and server for predicting damaging missense mutations. Nat Methods, 7(4), 248–249. 10.1038/nmeth0410-248

Ahlquist, R. P. (1948). A study of the adrenotropic receptors. Am J Physiol, 153(3), 586–600. 10.1152/ajplegacy.1948.153.3.586

Berman, H. M., Westbrook, J., Feng, Z., Gilliland, G., Bhat, T. N., Weissig, H., Shindyalov, I. N., & Bourne, P. E. (2000). The Protein Data Bank. Nucleic Acids Res, 28(1), 235–242. 10.1093/nar/28.1.235

Bertalan, E., Lesnik, S., Bren, U., & Bondar, A. N. (2020). Protein-water hydrogen-bond networks of G protein-coupled receptors: Graph-based analyses of static structures and molecular dynamics. J Struct Biol, 212(3), 107634. 10.1016/j.jsb.2020.107634

Boniecki, M., Rotkiewicz, P., Skolnick, J., & Kolinski, A. (2003). Protein fragment reconstruction using various modeling techniques. J Comput Aided Mol Des, 17(11), 725–738. 10.1023/b:jcam.0000017486.83645.a0

Brandes, N., Goldman, G., Wang, C. H., Ye, C. J., & Ntranos, V. (2023). Genome-wide prediction of disease variant effects with a deep protein language model. Nat Genet, 55(9), 1512–1522. 10.1038/s41588-023-01465-0

Chen, J. F., Sonsalla, P. K., Pedata, F., Melani, A., Domenici, M. R., Popoli, P., Geiger, J., Lopes, L. V., & de Mendonca, A. (2007). Adenosine A2A receptors and brain injury: broad spectrum of neuroprotection, multifaceted actions and “fine tuning” modulation. Prog Neurobiol, 83(5), 310–331. 10.1016/j.pneurobio.2007.09.002

Chen, T., & Guestrin, C. (2016). XGBoost: A Scalable Tree Boosting System Proceedings of the 22nd ACM SIGKDD International Conference on Knowledge Discovery and Data Mining, San Francisco, California, USA. 10.1145/2939672.2939785

Cheng, J., Novati, G., Pan, J., Bycroft, C., Zemgulyte, A., Applebaum, T., Pritzel, A., Wong, L. H., Zielinski, M., Sargeant, T., Schneider, R. G., Senior, A. W., Jumper, J., Hassabis, D., Kohli, P., & Avsec, Z. (2023). Accurate proteome-wide missense variant effect prediction with AlphaMissense. Science, 381(6664), eadg7492. 10.1126/science.adg7492

Choi, Y., & Chan, A. P. (2015). PROVEAN web server: a tool to predict the functional effect of amino acid substitutions and indels. Bioinformatics, 31(16), 2745–2747. 10.1093/bioinformatics/btv195

Christopoulos, A., May, L. T., Avlani, V. A., & Sexton, P. M. (2004). G-protein-coupled receptor allosterism: the promise and the problem(s). Biochem Soc Trans, 32(Pt 5), 873–877. 10.1042/BST0320873

Cline, M. S., & Karchin, R. (2011). Using bioinformatics to predict the functional impact of SNVs. Bioinformatics, 27(4), 441–448. 10.1093/bioinformatics/btq695

Collins, F. S., Guyer, M. S., & Charkravarti, A. (1997). Variations on a theme: cataloging human DNA sequence variation. Science, 278(5343), 1580–1581. 10.1126/science.278.5343.1580

de Araujo, A. D., Hoang, H. N., Lim, J., Mak, J. Y. W., & Fairlie, D. P. (2022). Tuning Electrostatic and Hydrophobic Surfaces of Aromatic Rings to Enhance Membrane Association and Cell Uptake of Peptides. Angew Chem Int Ed Engl, 61(29), e202203995. 10.1002/anie.202203995

Dong, C., Wei, P., Jian, X., Gibbs, R., Boerwinkle, E., Wang, K., & Liu, X. (2015). Comparison and integration of deleteriousness prediction methods for nonsynonymous SNVs in whole exome sequencing studies. Hum Mol Genet, 24(8), 2125–2137. 10.1093/hmg/ddu733

Drake, M. T., Shenoy, S. K., & Lefkowitz, R. J. (2006). Trafficking of G protein-coupled receptors. Circ Res, 99(6), 570–582. 10.1161/01.RES.0000242563.47507.ce

Duan, W., Gui, L., Zhou, Z., Liu, Y., Tian, H., Chen, J. F., & Zheng, J. (2009). Adenosine A2A receptor deficiency exacerbates white matter lesions and cognitive deficits induced by chronic cerebral hypoperfusion in mice. J Neurol Sci, 285(1-2), 39–45. 10.1016/j.jns.2009.05.010

Geurts, P., Ernst, D., & Wehenkel, L. (2006). Extremely randomized trees. Machine Learning, 63(1), 3–42. 10.1007/s10994-006-6226-1

Gromiha, M. M., & Ou, Y. Y. (2014). Bioinformatics approaches for functional annotation of membrane proteins. Brief Bioinform, 15(2), 155–168. 10.1093/bib/bbt015

Gudermann, T., Kalkbrenner, F., & Schultz, G. (1996). Diversity and selectivity of receptor-G protein interaction. Annu Rev Pharmacol Toxicol, 36, 429–459. 10.1146/annurev.pa.36.040196.002241

Hwang, S. K., & Kwon, S. (2015). Early-onset epileptic encephalopathies and the diagnostic approach to underlying causes. Korean J Pediatr, 58(11), 407–414. 10.3345/kjp.2015.58.11.407

Isberg, V., Mordalski, S., Munk, C., Rataj, K., Harpsoe, K., Hauser, A. S., Vroling, B., Bojarski, A. J., Vriend, G., & Gloriam, D. E. (2016). GPCRdb: an information system for G protein-coupled receptors. Nucleic Acids Res, 44(D1), D356–364. 10.1093/nar/gkv1178

Jumper, J., Evans, R., Pritzel, A., Green, T., Figurnov, M., Ronneberger, O., Tunyasuvunakool, K., Bates, R., Zidek, A., Potapenko, A., Bridgland, A., Meyer, C., Kohl, S. A. A., Ballard, A. J., Cowie, A., Romera-Paredes, B., Nikolov, S., Jain, R., Adler, J., Back, T., Petersen, S., Reiman, D., Clancy, E., Zielinski, M., Steinegger, M., Pacholska, M., Berghammer, T., Bodenstein, S., Silver, D., Vinyals, O., Senior, A. W., Kavukcuoglu, K., Kohli, P., & Hassabis, D. (2021). Highly accurate protein structure prediction with AlphaFold. Nature, 596(7873), 583–589. 10.1038/s41586-021-03819-2

Kawakami, N., Miyoshi, K., Horio, S., & Fukui, H. (2004). Beta(2)-adrenergic receptor-mediated histamine H(1) receptor down-regulation: another possible advantage of beta(2) agonists in asthmatic therapy. J Pharmacol Sci, 94(4), 449–458. 10.1254/jphs.94.449

Kawashima, S., Pokarowski, P., Pokarowska, M., Kolinski, A., Katayama, T., & Kanehisa, M. (2008). AAindex: amino acid index database, progress report 2008. Nucleic Acids Res, 36(Database issue), D202–205. 10.1093/nar/gkm998

Kulandaisamy, A., Binny Priya, S., Sakthivel, R., Tarnovskaya, S., Bizin, I., Honigschmid, P., Frishman, D., & Gromiha, M. M. (2018). MutHTP: mutations in human transmembrane proteins. Bioinformatics, 34(13), 2325–2326. 10.1093/bioinformatics/bty054

Kulandaisamy, A., Zaucha, J., Sakthivel, R., Frishman, D., & Michael Gromiha, M. (2020). Pred-MutHTP: Prediction of disease-causing and neutral mutations in human transmembrane proteins. Hum Mutat, 41(3), 581–590. 10.1002/humu.23961

Landrum, M. J., Lee, J. M., Riley, G. R., Jang, W., Rubinstein, W. S., Church, D. M., & Maglott, D. R. (2014). ClinVar: public archive of relationships among sequence variation and human phenotype. Nucleic Acids Res, 42(Database issue), D980–985. 10.1093/nar/gkt1113

Livesey, B. J., & Marsh, J. A. (2022). The properties of human disease mutations at protein interfaces. PLoS Comput Biol, 18(2), e1009858. 10.1371/journal.pcbi.1009858

Lundberg, S. M., & Lee, S.-I. (2017). A unified approach to interpreting model predictions Proceedings of the 31st International Conference on Neural Information Processing Systems, Long Beach, California, USA.

Luthy, R., McLachlan, A. D., & Eisenberg, D. (1991). Secondary structure-based profiles: use of structure-conserving scoring tables in searching protein sequence databases for structural similarities. Proteins, 10(3), 229–239. 10.1002/prot.340100307

Milligan, G., Ward, R. J., & Marsango, S. (2019). GPCR homo-oligomerization. Curr Opin Cell Biol, 57, 40–47. 10.1016/j.ceb.2018.10.007

Mooney, S. (2005). Bioinformatics approaches and resources for single nucleotide polymorphism functional analysis. Brief Bioinform, 6(1), 44–56. 10.1093/bib/6.1.44

Nathans, J., & Hogness, D. S. (1984). Isolation and nucleotide sequence of the gene encoding human rhodopsin. Proc Natl Acad Sci U S A, 81(15), 4851–4855. 10.1073/pnas.81.15.4851

Ng, P. C., & Henikoff, S. (2001). Predicting deleterious amino acid substitutions. Genome Res, 11(5), 863–874. 10.1101/gr.176601

Overington, J., Donnelly, D., Johnson, M. S., Sali, A., & Blundell, T. L. (1992). Environment-specific amino acid substitution tables: tertiary templates and prediction of protein folds. Protein Sci, 1(2), 216–226. 10.1002/pro.5560010203

Pandy-Szekeres, G., Caroli, J., Mamyrbekov, A., Kermani, A. A., Keseru, G. M., Kooistra, A. J., & Gloriam, D. E. (2023). GPCRdb in 2023: state-specific structure models using AlphaFold2 and new ligand resources. Nucleic Acids Res, 51(D1), D395–D402. 10.1093/nar/gkac1013

Park, P. S. (2019). Rhodopsin Oligomerization and Aggregation. J Membr Biol, 252(4-5), 413–423. 10.1007/s00232-019-00078-1

Pedregosa, F., Varoquaux, G., Gramfort, A., Michel, V., Thirion, B., Grisel, O., Blondel, M., Prettenhofer, P., Weiss, R., Dubourg, V., Vanderplas, J., Passos, A., Cournapeau, D., Brucher, M., Perrot, M., & Duchesnay, É. (2011). Scikit-learn: Machine Learning in Python. J. Mach. Learn. Res., 12(null), 2825–2830.

Popov, P., Bizin, I., Gromiha, M., A, K., & Frishman, D. (2019). Prediction of disease-associated mutations in the transmembrane regions of proteins with known 3D structure. PLoS One, 14(7), e0219452. 10.1371/journal.pone.0219452

Salon, J. A., Lodowski, D. T., & Palczewski, K. (2011). The significance of G protein-coupled receptor crystallography for drug discovery. Pharmacol Rev, 63(4), 901–937. 10.1124/pr.110.003350

Schipani, E., Kruse, K., & Juppner, H. (1995). A constitutively active mutant PTH-PTHrP receptor in Jansen-type metaphyseal chondrodysplasia. Science, 268(5207), 98–100. 10.1126/science.7701349

Schoneberg, T., & Liebscher, I. (2021). Mutations in G Protein-Coupled Receptors: Mechanisms, Pathophysiology and Potential Therapeutic Approaches. Pharmacol Rev, 73(1), 89–119. 10.1124/pharmrev.120.000011

Schoneberg, T., Schulz, A., Biebermann, H., Hermsdorf, T., Rompler, H., & Sangkuhl, K. (2004). Mutant G-protein-coupled receptors as a cause of human diseases. Pharmacol Ther, 104(3), 173–206. 10.1016/j.pharmthera.2004.08.008

Seifert, R., & Wenzel-Seifert, K. (2002). Constitutive activity of G-protein-coupled receptors: cause of disease and common property of wild-type receptors. Naunyn Schmiedebergs Arch Pharmacol, 366(5), 381–416. 10.1007/s00210-002-0588-0

Senese, N. B., Rasenick, M. M., & Traynor, J. R. (2018). The Role of G-proteins and G-protein Regulating Proteins in Depressive Disorders. Front Pharmacol, 9, 1289. 10.3389/fphar.2018.01289

Thompson, M. D., Percy, M. E., Cole, D. E. C., Bichet, D. G., Hauser, A. S., & Gorvin, C. M. (2024). G protein-coupled receptor (GPCR) gene variants and human genetic disease. Crit Rev Clin Lab Sci, 1–30. 10.1080/10408363.2023.2286606

Tin Kam, H. (1995, 14-16 Aug. 1995). Random decision forests. Proceedings of 3rd International Conference on Document Analysis and Recognition,

Ulloa-Aguirre, A., & Michael Conn, P. (2011). Pharmacoperones: a new therapeutic approach for diseases caused by misfolded G protein-coupled receptors. Recent Pat Endocr Metab Immune Drug Discov, 5(1), 13–24. 10.2174/187221411794351851

van der Westhuizen, E. T., Valant, C., Sexton, P. M., & Christopoulos, A. (2015). Endogenous allosteric modulators of G protein-coupled receptors. J Pharmacol Exp Ther, 353(2), 246–260. 10.1124/jpet.114.221606

Venkatakrishnan, A. J., Deupi, X., Lebon, G., Tate, C. G., Schertler, G. F., & Babu, M. M. (2013). Molecular signatures of G-protein-coupled receptors. Nature, 494(7436), 185–194. 10.1038/nature11896

Venkatakrishnan, A. J., Ma, A. K., Fonseca, R., Latorraca, N. R., Kelly, B., Betz, R. M., Asawa, C., Kobilka, B. K., & Dror, R. O. (2019). Diverse GPCRs exhibit conserved water networks for stabilization and activation. Proc Natl Acad Sci U S A, 116(8), 3288–3293. 10.1073/pnas.1809251116

Wang, D., Li, J., Wang, E., & Wang, Y. (2024). DVA: predicting the functional impact of single nucleotide missense variants. BMC Bioinformatics, 25(Suppl 1), 100. 10.1186/s12859-024-05709-6

Weis, W. I., & Kobilka, B. K. (2018). The Molecular Basis of G Protein-Coupled Receptor Activation. Annu Rev Biochem, 87, 897–919. 10.1146/annurev-biochem-060614-033910

Wootten, D., Christopoulos, A., Marti-Solano, M., Babu, M. M., & Sexton, P. M. (2018). Mechanisms of signalling and biased agonism in G protein-coupled receptors. Nat Rev Mol Cell Biol, 19(10), 638–653. 10.1038/s41580-018-0049-3

Xiong, D., Lee, D., Li, L., Zhao, Q., & Yu, H. (2022). Implications of disease-related mutations at protein-protein interfaces. Curr Opin Struct Biol, 72, 219–225. 10.1016/j.sbi.2021.11.012

Zalewska, M., Siara, M., & Sajewicz, W. (2014). G protein-coupled receptors: abnormalities in signal transmission, disease states and pharmacotherapy. Acta Pol Pharm, 71(2), 229–243. https://www.ncbi.nlm.nih.gov/pubmed/25272642

