## Supplementary Material for "From Mutations to Disease: Computational Analysis and Interpretation of GPCR-Associated Pathogenicity"

### SUPPLEMENTARY MATERIAL AND METHODS

#### Evaluation metrics for classification models

#### Accuracy

Accuracy is a measure that gauges the overall correctness of a classification model by considering the ratio of correctly predicted instances to the total instances. The accuracy formula is expressed as:

$$\text{Accuracy} = \frac{TP + TN}{TP + TN + FP + FN}$$

#### Matthew's correlation coefficient (MCC)

MCC is a crucial metric in assessing classification models, especially when dealing with imbalanced datasets. MCC considers both false and true positive and negative predictions to offer a comprehensive evaluation of imbalanced learning. Functioning as a correlation coefficient, MCC ranges from -1 to +1. A score of +1 indicates a perfect positive correlation, while -1 signifies an inverse correlation between the predictions and the true class labels. Meanwhile, a score of 0 suggests a performance equivalent to a random classifier. MCC is determined in the following equation:

$$\text{MCC} = \frac{\text{TP} \times \text{TN} - \text{FP} \times \text{FN}}{\sqrt{(\text{TP}+\text{FP}) \times (\text{FN}+\text{TN}) \times (\text{TP}+\text{FN}) \times (\text{FP}+\text{TN})}}$$

Here, TP stands for True Positives, TN for True Negatives, FP for False Positives, and FN for False Negatives.

#### F1 Score

The F1-score is a metric that combines precision and recall, considering both false positives and false negatives, with a focus on imbalanced datasets. It calculates the harmonic mean of precision and recall, assigning different weights to each class based on their prevalence in the dataset. The formula for the F1 Score is given by:

$$\text{F1 Score} = \frac{TP}{TP + \frac{1}{2}(FP + FN)}$$

**Weighted F1 Score**. Here, we employed a weighted version of the F1 score, presented in the following equation. In this equation, $N$ represents the number of classes, $F1\,Score_i$ is the F1-score for the class $i$ and $Wi$ is the weight assigned to the class $i$, usually determined by the class distribution in the dataset. The Weighted F1 Score provides a more nuanced evaluation of a model's performance, particularly in scenarios where class imbalances exist.

$$\text{Weighted F1 Score} = \sum_{i=1}^{N} w_i \times \text{F1 Score}_i$$

**Area under the curve score (AUC)**

A ROC curve, short for the Receiver Operating Characteristic curve, is a graphical representation that illustrates the performance of a binary classifier system as its discrimination threshold is varied. It is created by plotting the fraction of true positives out of the positives (TPR = true positive rate) vs. the fraction of false positives out of the negatives (FPR = false positive rate), at various threshold settings. TPR is also known as sensitivity, and FPR is one minus the specificity or true negative rate.

The AUC computed in this study, considers the area under the ROC curve. By doing so, the curve information is summarised in one number.

**Sensitivity**

This metric is equivalent to the true positive rate. It is the probability of a positive test result, conditioned on the individual truly being positive.

$$\begin{aligned}\text{sensitivity} &= \frac{\text{number of true positives}}{\text{number of true positives} + \text{number of false negatives}}\\ &= \frac{\text{number of true positives}}{\text{total number of sick individuals in population}}\\ &= \text{probability of a positive test given that the patient has the disease}\end{aligned}$$

**Specificity**

This metric is equivalent to the true negative rate. It is the probability of a negative test result, conditioned on the individual truly being negative.

$$\text{specificity} = \frac{\text{number of true negatives}}{\text{number of true negatives} + \text{number of false positives}}$$

$$= \frac{\text{number of true negatives}}{\text{total number of well individuals in population}}$$

$$= \text{probability of a negative test given that the patient is well}$$

### SUPPLEMENTARY RESULTS

#### MutHTP data

#### Benchmark, analysis and understanding of predictive mutagenesis tools on GPCR mutations

The results obtained from the evaluation of tools for the MutHTP dataset, as detailed in Table S3 (Figure S8), diverged significantly from those observed in the ClinVar dataset. The performances were considerably lower, with the highest MCC reaching only 0.13 for AlphaMissense. While SIFT, PolyPhen-2, and PROVEAN exhibited similar performances, with accuracies of 0.67, 0.70, and 0.68 respectively, AlphaMissense demonstrated distinct characteristics with lower accuracy and F1-score weighted compared to the other tools. The low classification accuracy displayed by all tools, pointed out that these tools are unreliable when categorising GPCR variants as pathogenic or benign for the MutHTP data set. This inconsistency highlights the importance of dataset-specific evaluation and the need for further investigation to improve predictive accuracy and reliability in mutation classification. As aforementioned, all evaluation metrics for classification are also detailed in the Supplementary Materials.

Following the same principle for the previous database, we also generated a heatmap Figure S8 to provide a visual comparison of tools' accuracy on the MutHTP dataset. As observed in our benchmark (Table S3), PolyPhen-2 and SIFT presented very similar performance, with PolyPhen-2 being slightly better, with an accuracy of 0.70 while SIFT displayed an accuracy of 0.67. Nevertheless, the performance, considering all tools, was much lower when compared to the performance observed in the ClinVar dataset (see Table 1). The MCC was as low as 0.09 for PROVEAN, and ESM1b, and as high as 0.11 for PolyPhen-2 and AlphaMissense. These results differ strongly to ClinVar, where the lowest MCC was 0.50 for PolyPhen-2. A possible cause of this is the fact that MutHTP is heavily based on mutations related to cancer, which have a higher level of complexity. Predicting the effects of mutations that cause cancer is particularly challenging due to the intrinsic combinations of multiple factors that lead to cancer. Cancer development involves intricate biological pathways where mutations can disrupt numerous signalling cascades, leading to uncontrolled cell growth and metastasis (Hanahan & Weinberg, 2011). The tumour microenvironment adds another layer of complexity. For instance, it consists of a dynamic milieu of cancer cells, stromal cells, immune cells, and extracellular matrix components that interact and evolve, significantly influencing tumour progression and therapeutic response (Quail & Joyce, 2013). Furthermore, distinguishing between driver mutations, which directly contribute to cancer development, and passenger mutations, which are incidental by-products of the cancer genome's instability, complicates the identification of clinically relevant mutations (Vogelstein et al., 2013). This multifaceted nature of cancer biology contributes to the difficulty in accurately predicting the pathogenic potential of specific mutations.

Additionally, we also evaluated the overall performance of our predictors on cases where the PolyPhen-2 predictor made incorrect predictions (benign or pathogenic). This analysis aimed to determine whether other predictors could compensate for these errors. This analysis is represented in a second heatmap Figure S9 that includes only GPCR mutations where PolyPhen-2 failed to correctly identify the clinical significance of the variant. It is possible to observe that potentially the combination of PolyPhen-2, PROVEAN and SIFT could support a slightly better performance (more blue lines on PROVEAN and SIFT when comparing them to the other tools, which indicates correct predictions). Nevertheless, caution would be highly advisable considering the previously mentioned low performance of all of them.

To further investigate this scenario, we also generated confusion matrices to provide a detailed view of how each predictor performs in terms of correctly and incorrectly classifying the clinical significance of variants that PolyPhen-2 failed to predict properly (Figure S10). For instance, a higher count of true benign (top-left cell) and true pathogenic (bottom-right cell) cells indicates better performance, while higher counts in the false pathogenic (top-right cell) and false benign (bottom-left cell) cells highlight areas where the predictor struggles. Considering these premises, it is possible to observe that indeed PROVEAN (Figure S10C) and SIFT (Figure S10D) had a better performance when considering the number of true pathogenic. PROVEAN has a count of 355 and SIFT has a count of 276 of true pathogenic. AlphaMissense, and ESM1b, presented slightly better true benign counts, 118 and 82 respectively. Still, all tools rendered a high amount of false benign (i.e., pathogenic mutations being missed by the predictor), which is very alarming when dealing with predictions of mutations leading to diseases. One suggestion would be to apply more weight to the pathogenic class during the predictions. This weighting scheme could change the probability of classification, possibly rendering a better predictive performance on classifying GPCR mutations into benign and pathogenic.

Subsequently, we also analysed the performance of each predictor considering each one of the following types of information about the mutations presented in the MutHTP dataset: UniProt ID, GPCRs classes, phenotype, and cellular position.

When considering the grouping by UniProt ID (Figure S11), a similar pattern of high variability in accuracy is observed, as was seen with the ClinVar data. For instance, the accuracy of the AlphaMissense tool can be as low as 35% for predicting mutations in the receptor UniProt ID Q969V1 (Melanin-concentrating hormone receptor 2). In contrast, the PolyPhen-2 tool achieves an accuracy as high as 93% for the GPCR with UniProt ID P30518 (Vasopressin V2 receptor). These findings point to the significant variability in tool performance depending on the specific receptor being analysed. Notably, the Vasopressin V2 receptor was also highlighted during the evaluation of the ClinVar dataset. As expected, the accuracy levels for this specific receptor were similar across all tools on both data sets (ClinVar and MutHTP).

The histogram obtained from the analysis by grouping the mutations according to GPCR classes (Figure S12) revealed that the predictions for Class C (Metabotropic glutamate) GPCRs using MutHTP were more accurate. Conversely, predictions for Class A (Rhodopsin-like) GPCRs were the least accurate across nearly all tools. As low as 41% and 49% for AlphaMissense, and ESM1b, respectively. This trend reinforces that current predictive tools may not be optimally designed for Rhodopsin-like (Class A) GPCRs. This consistent pattern suggests a need for further refinement and specialisation of these tools to improve their performance for all classes. There is a need for specialisation of predictive tools for GPCR's Class A, which represents the classes with more GPCRs in nature.

Subsequently, we evaluated the precision of the tools by grouping mutations according to their phenotype (Figure S13) as described in the MutHTP dataset. The bar plot illustrates the accuracy of different predictive tools in identifying the pathogenicity of mutations across various phenotypic disease categories. Each bar represents the percentage of correct predictions made by a specific predictor, with different colours denoting the phenotypic disease categories. Like our observations with the ClinVar data, from the plot, it is evident that the prediction accuracies vary significantly across different predictors and phenotypes. For instance, the tool PolyPhen-2 demonstrates the highest accuracy, exceeding 90%, across several phenotypes, including Congenital Disorders Of Metabolism, Endocrine And Metabolic Diseases, and Urinary System Diseases. AlphaMissense demonstrated the lowest accuracy on average. Demonstrating accuracies as low as 38% for cancer-related mutations, and a maximum of 80% for mutations related to Nervous System diseases. For all predictors, the accuracy-related to cancer mutations is the lowest when compared to other phenotypes. This result reinforces the assumption that the predictive performance is much lower in the MutHTP dataset compared to the ClinVar dataset because MutHTP is heavily based on mutations related to cancer. This variability highlights the differing capabilities of these tools in handling specific disease categories, indicating that some tools are better tailored for certain phenotypes over others. Furthermore, it illustrates the challenges of predicting mutations leading to cancer.

At last, we evaluated the tools' accuracy according to cellular localisation of the mutation (transmembrane, cytoplasmic domain, and non-cytoplasmic domain) (Figure S14). It is observed that PolyPhen-2 and PROVEAN rank as the best performers across all localisations. Considering cytoplasmic domain mutations, the accuracies were the highest seen in all cases, 71% and 73%, for PROVEAN and PolyPhen-2, respectively. Conversely, AlphaMissense consistently shows the lowest performance, especially in the cytoplasmic domain (i.e., 36%). The non-cytoplasmic domain poses the greatest challenge, with lower accuracy across almost all tools. These trends highlight the significance of the physical context of mutations in prediction accuracy, indicating that enhancements in predictive models should focus on improving accuracy in non-cytoplasmic domains. This chart emphasises the need for localisation-specific training and refinement in predictive tools to improve overall performance. These results emphasise the importance of considering characteristics-specific nuances in mutation prediction and highlight areas for further tool refinement.

According to the results, predicting the effects of GPCR mutations leading to cancer is complex and therefore challenging to be computationally identified by machine learning pipelines and models. We assume that this is caused by the various factors involving cancer development, such as the involvement of complex biological pathways, the influence of the tumour microenvironment, and the need to distinguish mutations that are really involved and mutations that are present but have no impact on cancer development. Given these challenges, the development of specialised tools is of major importance. As seen through these results, new tools should consider information regarding the mutations, such as subcellular localisation, class of the GPCR, the specific GPCR involved in the disease, and disease phenotypes caused by the mutation. These predictive tools may ultimately contribute to more effective diagnostics and personalised treatment strategies for cancer patients, especially if their results are combined or redesigned to consider GPCR's inherent characteristics.

**Statistical analysis and interpretation of GPCR mutations**

Next, we aimed to identify features, or molecular drivers, that can distinguish between pathogenic and benign GPCR mutations within the MutHTP dataset. Descriptive features showing a high correlation with outcomes were explored in detail to provide insights into the molecular mechanisms that lead to disease development. Similarly to the analysis made in the previous section, our approach involved statistical analysis of a diverse array of features to uncover the molecular drivers of mutations associated with GPCR-related disorders. We evaluated the predictive potential of these features in mutation classification using machine learning models (see the Methodology section), and Shapley Additive Explanations (SHAP). When analysing the impact of deleterious mutations in GPCRs using the MutHTP dataset, we found that all features exhibited a notably low screening performance. This agrees with the performance observed when assessing the available predictive tools. Furthermore, the results across pathogenic and benign mutations showed significant similarities (see Figure S15 and Table S2), underlining the challenge of accurately classifying mutations in the MutHTP dataset.

When evaluating the SHAP values (see Figure S15) for the MutHTP dataset, the three most important features were DOSZ010101 (extracted from AAindex, (Kawashima et al., 2008), using an Amino acid similarity matrix based on the sausage force field, (Dosztanyi & Torda, 2001)), Hbond (number of hydrogen bonds on the mutation site), and the graph-based signature Acc:Hydro-3.00 (this feature represents the quantification of pairs of pharmacophoric regions within a defined distance threshold surrounding the mutation site. In this case, it indicates the count of pairs of Hydrogen bond acceptors atoms and hydrophobic atoms within a maximum distance of 3 angstroms from the mutation site). According to the violin plots (Figure S16 A, B, and C), for all three features, the distribution of values is very similar (first quartile, median, third quartile). The mentioned distribution makes it hard to draw conclusions about them. However, it highlights the complexities in classifying mutations from the MutHTP dataset. Similarly to the violin plots, the patterns on the SHAP plot (see Figure S15) for the features DOSZ010101, Hbond, and Acc:Hydro-3.00 are mixed to a great extent, with higher and low values are correlated to both pathogenic and benign, indicating high difficulty in distinguishing between pathogenic and benign mutations in MutHTP dataset with well-known and robust sequence- and structural descriptors.

**SUPPLEMENTARY TABLES**

**Table S1: Evaluated features for ClinVar dataset.**

|  | Feature ID | Description | Reference |
|---|---|---|---|
| 1 | sift_pred | SIFT predictions | Ng and Henikoff (2001) |
| 2 | pph2_pred | PolyPhen-2 predictions | Adzhubei et al. (2010) |
| 3 | LUTR910101 | AAindex: Structure-based comparison table for outside other class | Luthy et al. (1991) |
| 4 | flu_reach | Normal Modes Analysis: Numeric vector of atomic fluctuations using 'REACH' force field (Bio3D package) | Skjaerven et al. (2014) |
| 5 | OVEJ920100_RSA | AAindex: OVEJ920105. Environment-specific amino acid substitution matrix for inaccessible residues | Overington et al. (1992) |
| 6 | Proximal | Structure-based, Arpeggio: Denotes if the atom is > the VdW interaction distance, but within 5 Angstroms of other atom(s). | Jubb et al. (2017) |
| 7 | Aro:Hydro-6.00 | Graph-based signature: Presence of pharmacophoric pairs Aromatic: Hydrophobic in a distance cut-off of 6 angstroms. | Pires et al. (2014) |
| 8 | gpcr_class | Class of the GPCR (A, B, C, D, F) | Nil |
| 9 | BONM030102 | AAindex: Quasichemical statistical potential for the intermediate orientation of interacting side groups | Boniecki et al. (2003) |
| 10 | Aro:Hydro-5.00 | Graph-based signature: Presence of pharmacophoric pairs Aromatic: Hydrophobic in a distance cut-off of 5 angstroms. | Pires et al. (2014) |
| 11 | Don:Sul-5.50 | Graph-based signature: Presence of pharmacophoric pairs Hydrogen bond Donor: Sulphur in a distance cut-off of 5.5 angstroms. | Pires et al. (2014) |
| 12 | non_cytosol | Mutation occurring outside the cytosol. | Nil |
| 13 | Neg:Neutral-1.00 | Graph-based signature: Presence of pharmacophoric pairs Negative: Neutral in a distance cut-off of 1 angstrom. | Pires et al. (2014) |

| | | | |
|---|---|---|---|
| 14 | Hydro:Pos-1.50 | Graph-based signature: Presence of pharmacophoric pairs Hydrophobic: Positive in a distance cut-off of 1.5 angstroms. | Pires et al. (2014) |

**Table S2: Evaluated features for MutHTP dataset.**

| | Feature ID | Description | Reference |
|---|---|---|---|
| 1 | DOSZ010101 | Amino acid similarity matrix based on the sausage force field | Dosztanyi and Torda (2001) |
| 2 | Hbond | Structure-based, Arpeggio: Denotes if the atom forms a hydrogen bond. | Jubb et al. (2017) |
| 3 | Acc:Hydro-3.00 | Graph-based signature: Presence of pharmacophoric pairs Hydrogen bond acceptor: Hydrophobic in a distance cut-off of 3 angstroms. | Pires et al. (2014) |
| 4 | Neg:Neg-6.00 | Graph-based signature: Presence of pharmacophoric pairs Negative: Negative in a distance cut-off of 6 angstroms. | Pires et al. (2014) |
| 5 | MetalSulphur-PI | Structure-based, Arpeggio: Methionine sulphur - π ring interaction | Jubb et al. (2017) |
| 6 | d_Carbonyl | Structure-based, Arpeggio: Denotes differences due to the mutation on carbonyl-carbon:carbonyl-carbon interactions. | Jubb et al. (2017) |
| 7 | Donor-PI | Structure-based, Arpeggio: Hydrogen Bond donor - π interactions | Jubb et al. (2017) |
| 8 | FromGLY | Mutations from Glycine | Nil |
| 9 | Aromatic | Structure-based, Arpeggio: Denotes an aromatic ring atom interacting with another aromatic ring atom. | Jubb et al. (2017) |
| 10 | Neg | Number of negative atoms | Pires et al. (2014) |
| 11 | Neg:Pos-5.00 | Graph-based signature: Presence of pharmacophoric pairs Negative: Positive in a distance cut-off of 5 angstroms. | Pires et al. (2014) |

| | | | |
|---|---|---|---|
| 12 | Neg:Pos-2.50 | Graph-based signature: Presence of pharmacophoric pairs Negative: Positive in a distance cut-off of 2.5 angstroms. | Pires et al. (2014) |

**Table S3: Predictive performance of mutation predictive tools for assessing the impact of GPCR mutations derived from MutHTP.**

| Method | Accuracy | MCC | F1-score weighted | AUC | Sensitivity | Specificity |
|---|---|---|---|---|---|---|
| **AlphaMissense** | 0.45 | 0.11 | 0.55 | 0.61 | 0.42 | 0.79 |
| **SIFT** | 0.67 | 0.10 | 0.75 | 0.59 | 0.68 | 0.49 |
| **PolyPhen-2** | 0.70 | 0.11 | 0.77 | 0.59 | 0.72 | 0.46 |
| **ESM1b (used the threshold of −7.5, lower than −7.5means the mutation is pathogenic) (https://doi.org/10.1038/s41588-023-01465-0)** | 0.54 | 0.09 | 0.64 | 0.59 | 0.53 | 0.65 |
| **PROVEAN (considered a score higher than -2.5 to be benign and lower than or equal to -2.5 to be pathogenic). For 40 mutations generating PROVEAN scores failed.** | 0.68 | 0.09 | 0.75 | 0.58 | 0.69 | 0.47 |

### SUPPLEMENTARY FIGURES

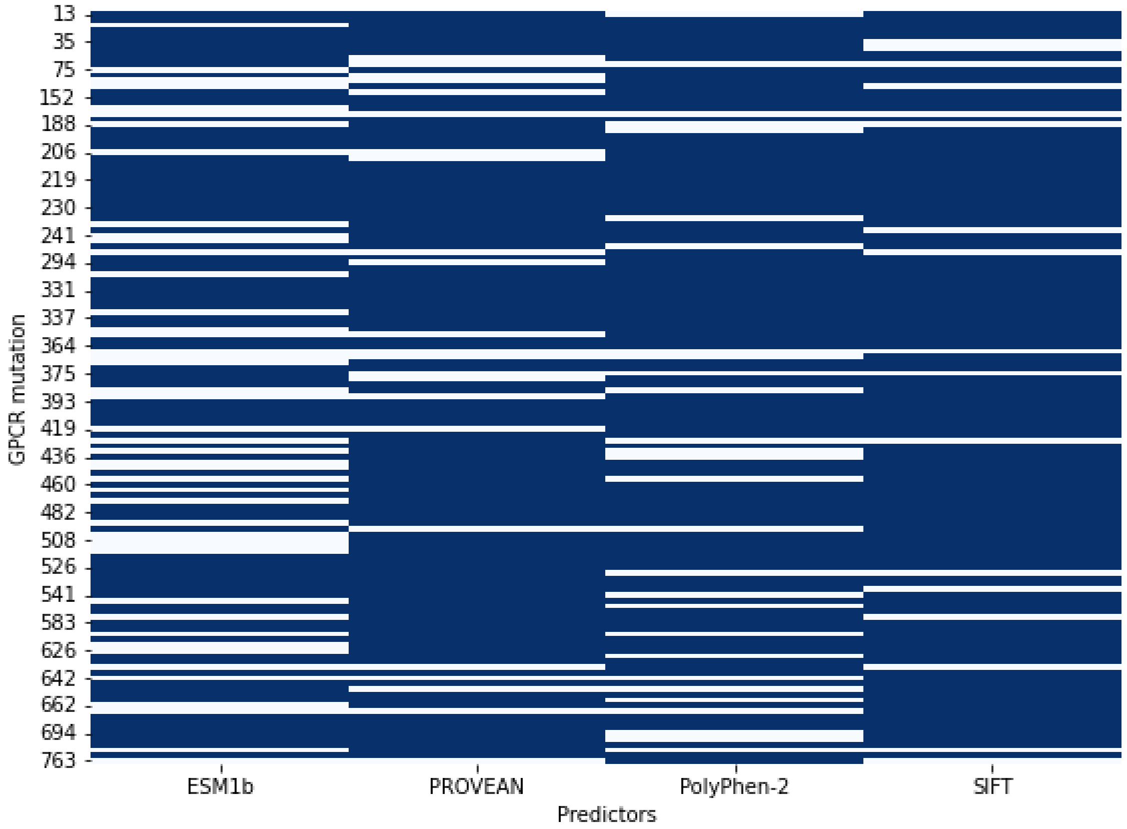


**Figure S1: Heatmap showing the accuracy of predictors in cases where the AlphaMissense predictor made an incorrect prediction. Each row represents a GPCR mutation in the ClinVar data set where AlphaMissense was wrong, and each column corresponds to the other predictors. A blue cell indicates a correct prediction by the corresponding predictor for that GPCR mutation. White cells, in turn, indicate the predictor's classification errors.**

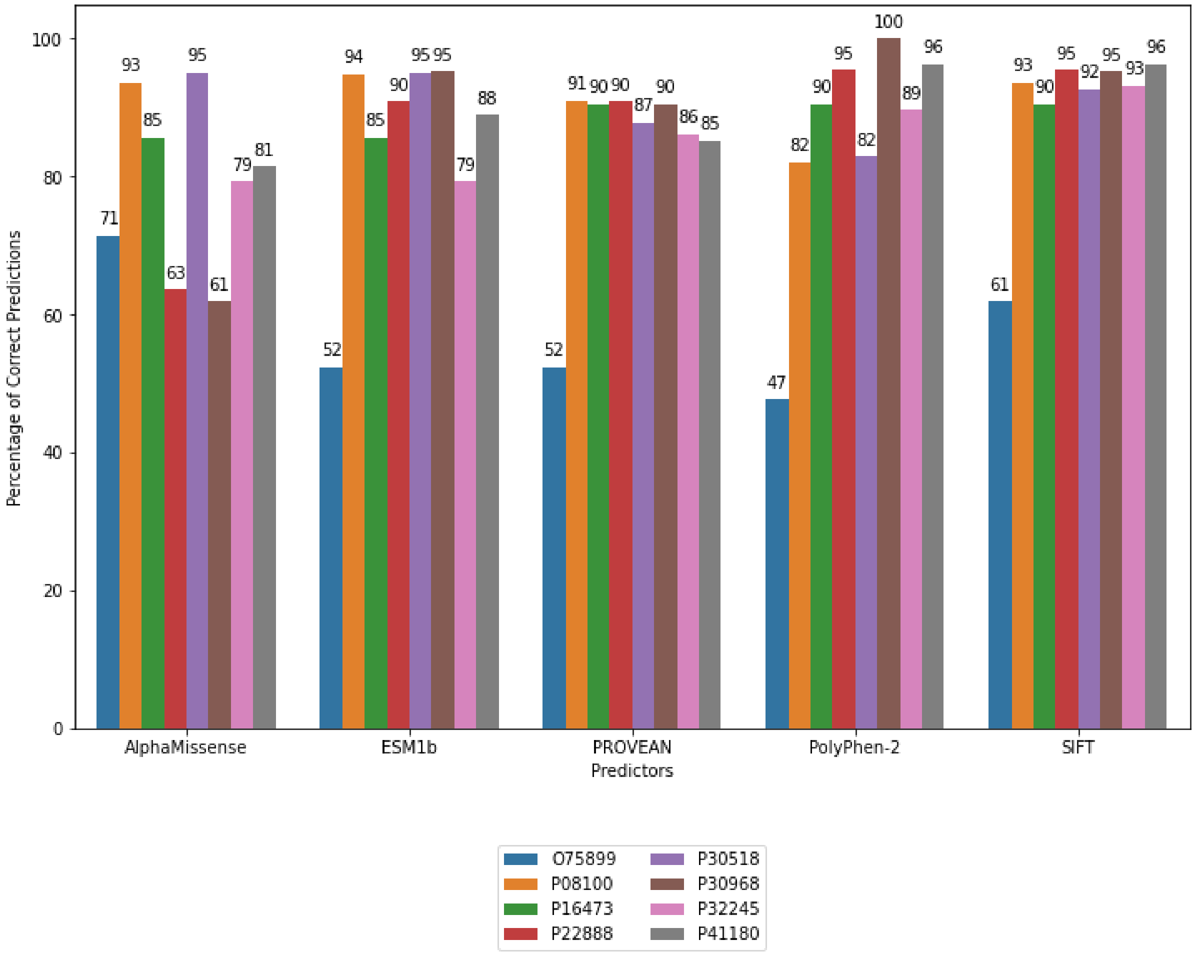


**Figure S2: Bar plot showing the percentage of correct predictions by predictor and UniProt, filtered to include only groups with more than 20 total cases (ClinVar data set).**

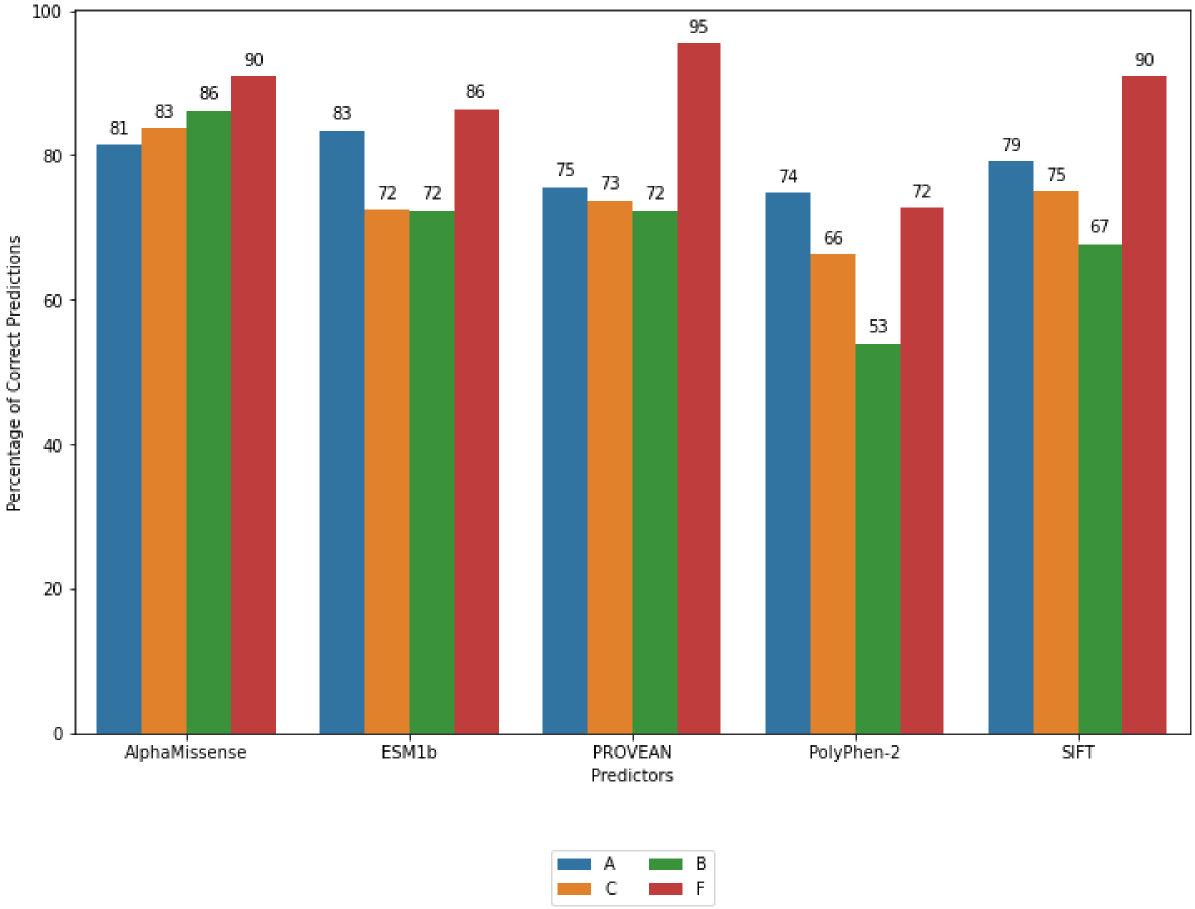


**Figure S3: Bar plot showing the percentage of correct predictions by predictor and UniProt (ClinVar data set).**

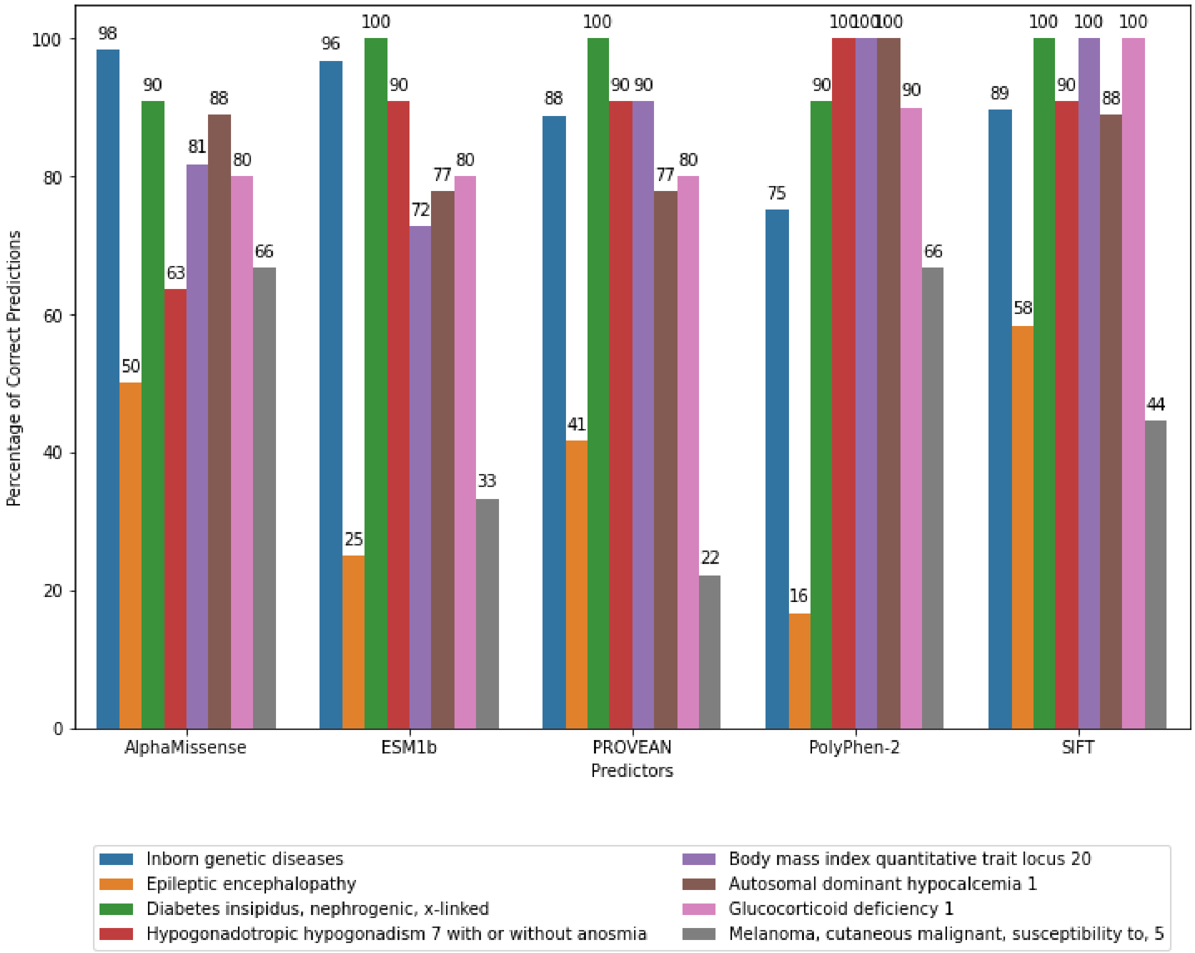


**Figure S4: Bar plot showing the percentage of correct predictions by predictor and disease phenotype, filtered to include only groups with more than 8 total cases (ClinVar data set).**

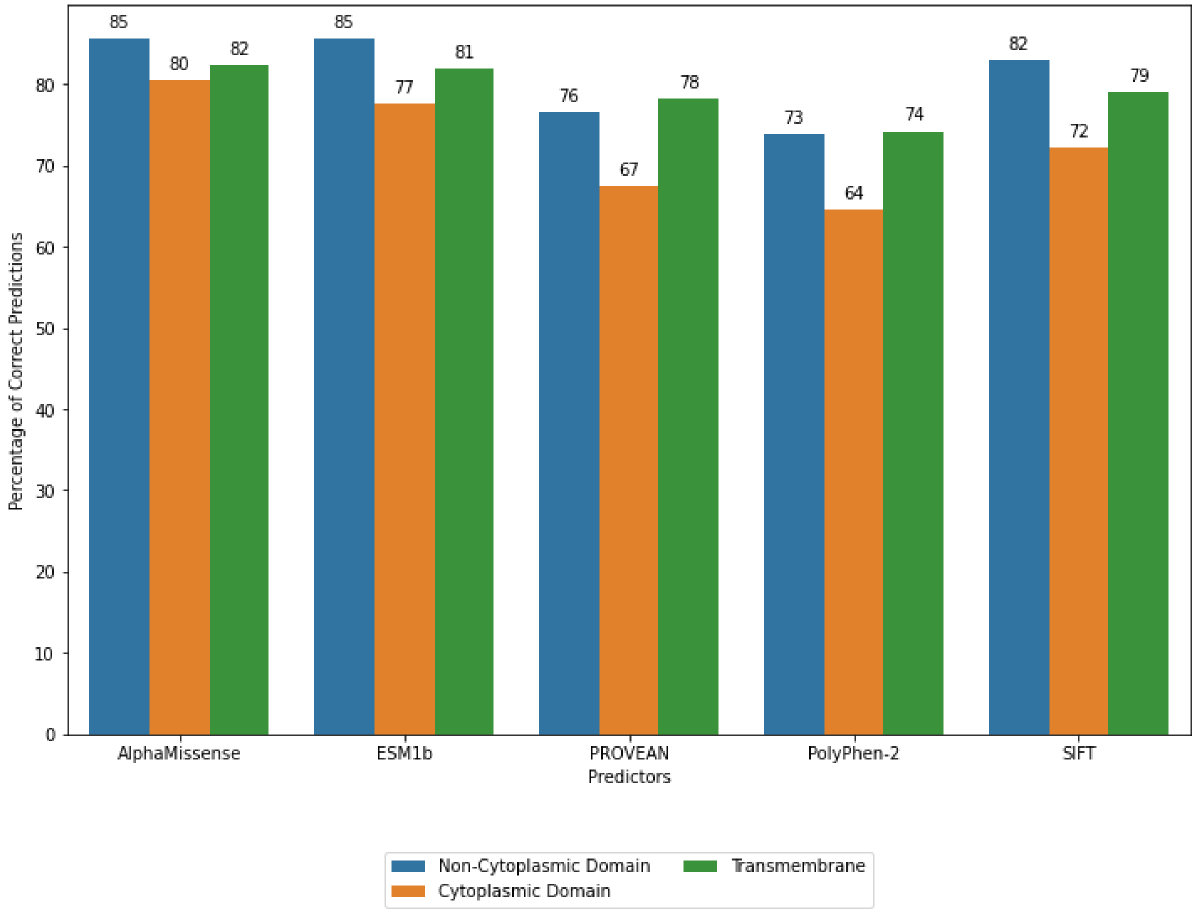


**Figure S5: Bar plot showing the number of correct predictions by predictor and cellular localisation (ClinVar data set).**

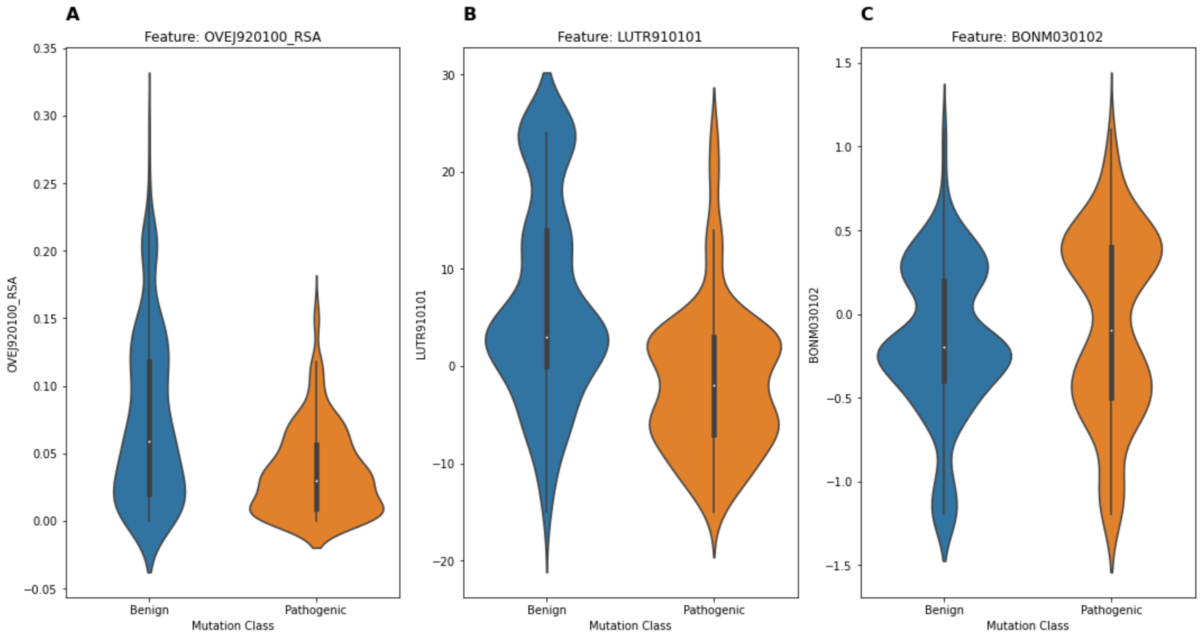


**Figure S6: Violin plot representing the best three AAindex features when analysing the ClinVar dataset. The violin plot displays the distribution of the three features (A: OVEJ920100_RSA, B: LUTR910101, and C: BONM030102) values for mutation classes. Pathogenic mutations are represented by the colour orange, while benign mutations**

**are represented by the colour blue. The width of the plot at each value indicates the density of data points.**

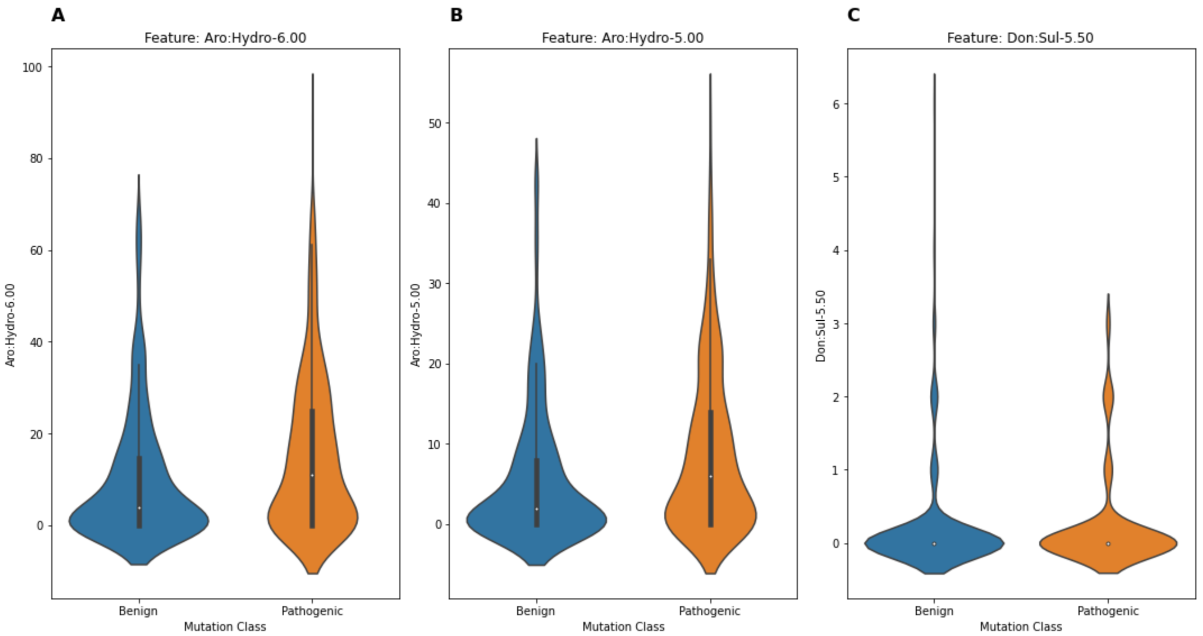


**Figure S7: Violin plot representing the best 3 graph-based signatures when analysing the ClinVar dataset. The violin plot displays the distribution of the 3 features (A: Aro:Hydro-6.00, B: Aro:Hydro-5.00, C: Don:Sul-5.50) values for mutation classes. Pathogenic mutations are represented by orange, while benign mutations are represented by blue. The width of the plot at each value indicates the density of data points.**

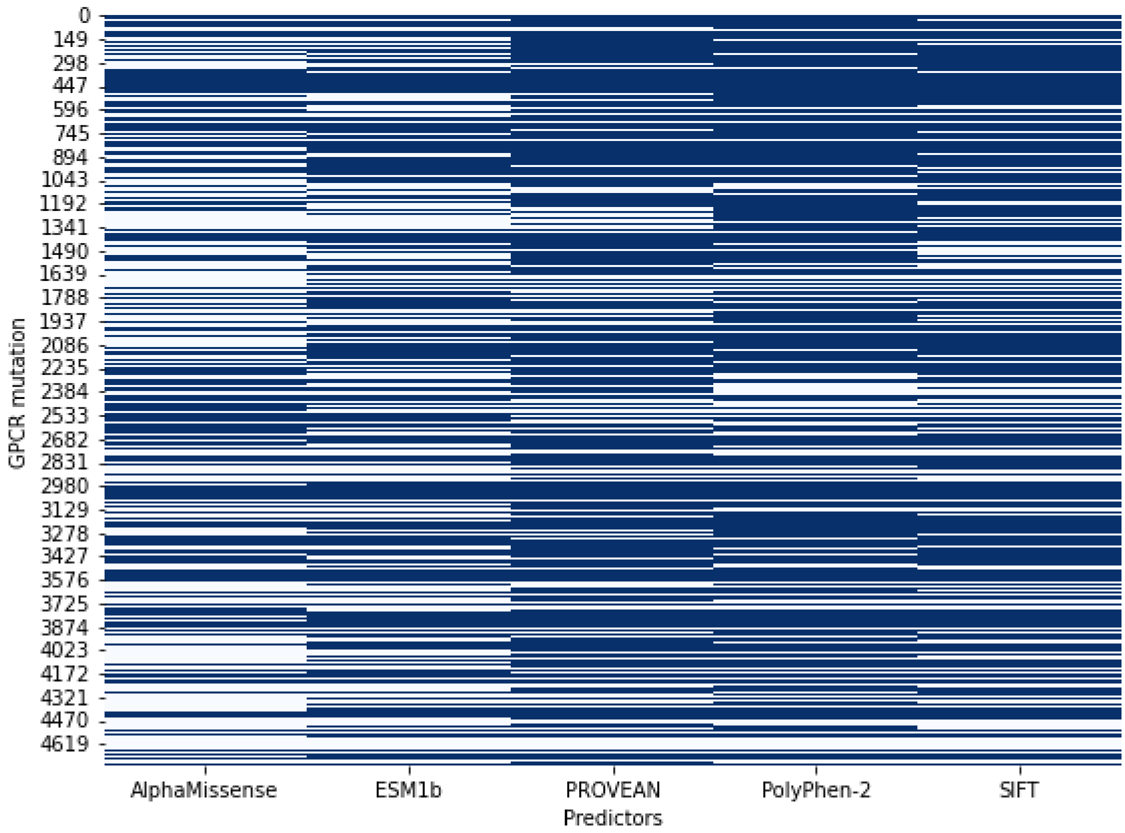


**Figure S8: Heatmap depicting the accuracy of various predictors in identifying the clinical significance of variants. Each row in the heatmap represents a distinct GPCR mutation in the MutHTP data set, while each column corresponds to a specific predictor used in the analysis. Blue cells indicate instances where the predictor correctly identified the clinical significance of the variant for that GPCR mutation. White cells, in turn, indicate the predictor's classification errors. This visual representation allows for a comprehensive comparison of the performance of different predictors across multiple GPCR mutations.**

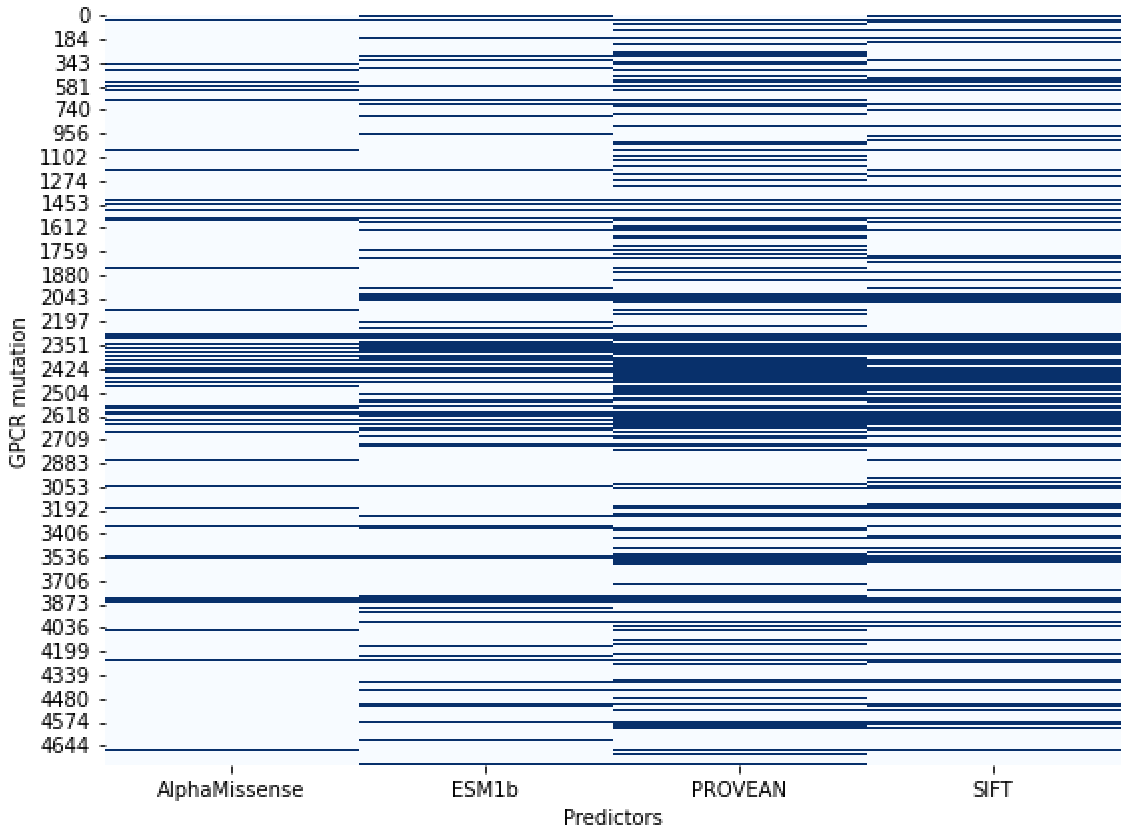


**Figure S9: Heatmap showing the accuracy of predictors in cases where the PolyPhen-2 predictor made an incorrect prediction. Each row represents a GPCR mutation in the MutHTP data set where the PolyPhen-2 prediction was wrong, and each column corresponds to the other predictors. A blue cell indicates a correct prediction by the corresponding predictor for that GPCR mutation.**

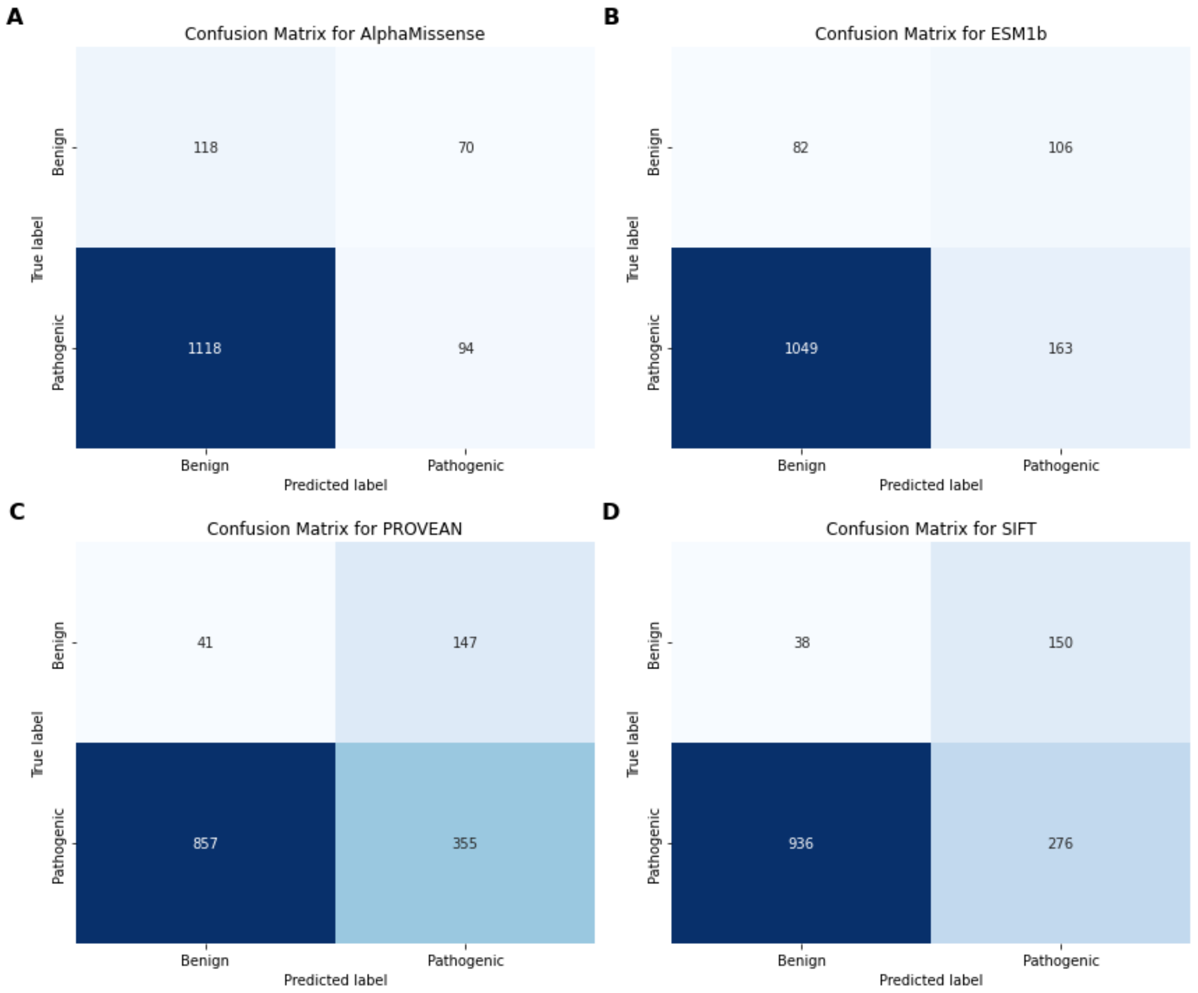


**Figure S10: Confusion matrices (MutHTP data set) for the A: AlphaMissense, B: ESM1b, C: PROVEAN, and D: 'SIFT' predictors, showing their performance in classifying the clinical significance of variants. Each subplot presents a matrix with true labels on the y-axis and predicted labels on the x-axis. The cell values indicate the number of GPCR mutations in each category. These matrices illustrate the accuracy of each predictor, with a focus on cases where PolyPhen-2 made incorrect predictions.**

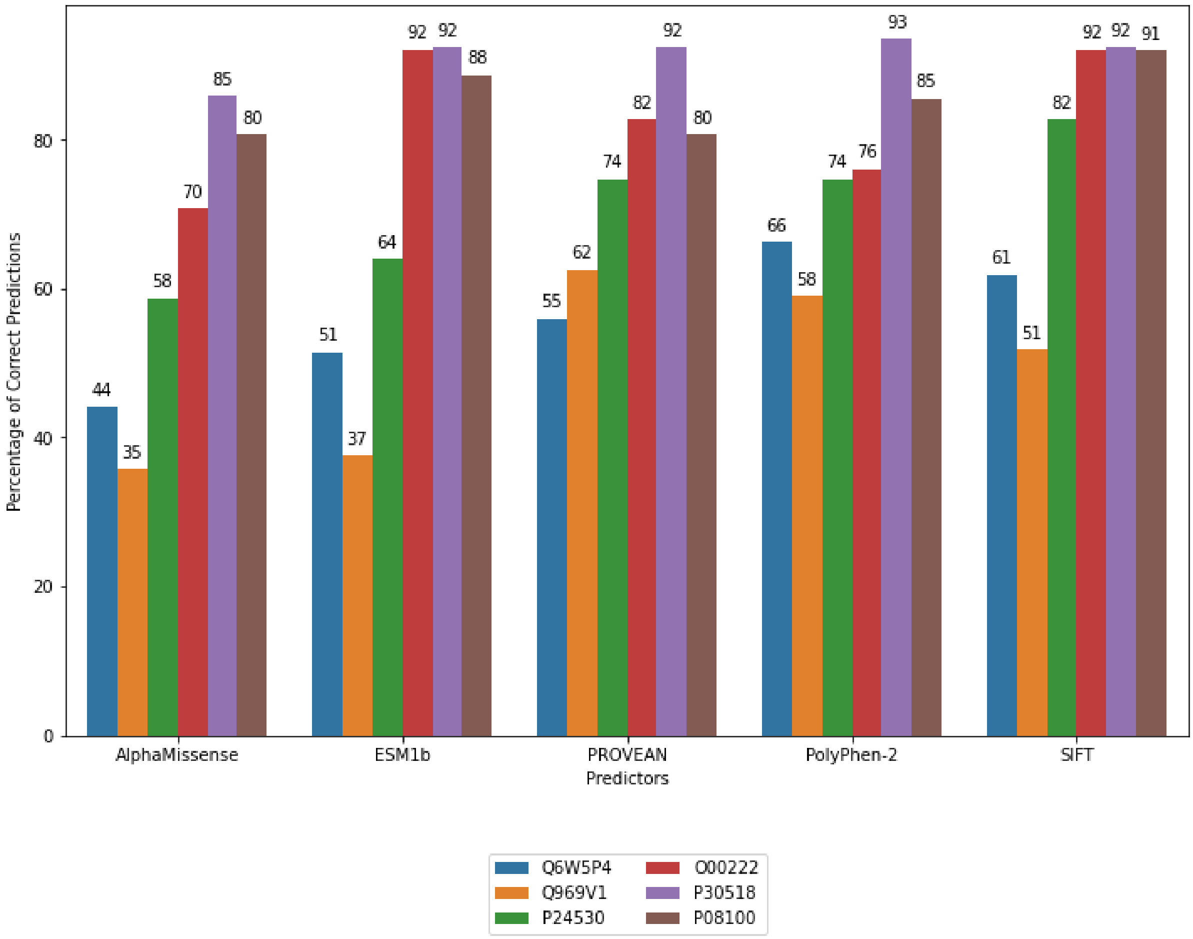


**Figure S11: Bar plot showing the number of correct predictions by predictor and UniProt, filtered to include only groups with more than 55 total cases (MutHTP data set).**

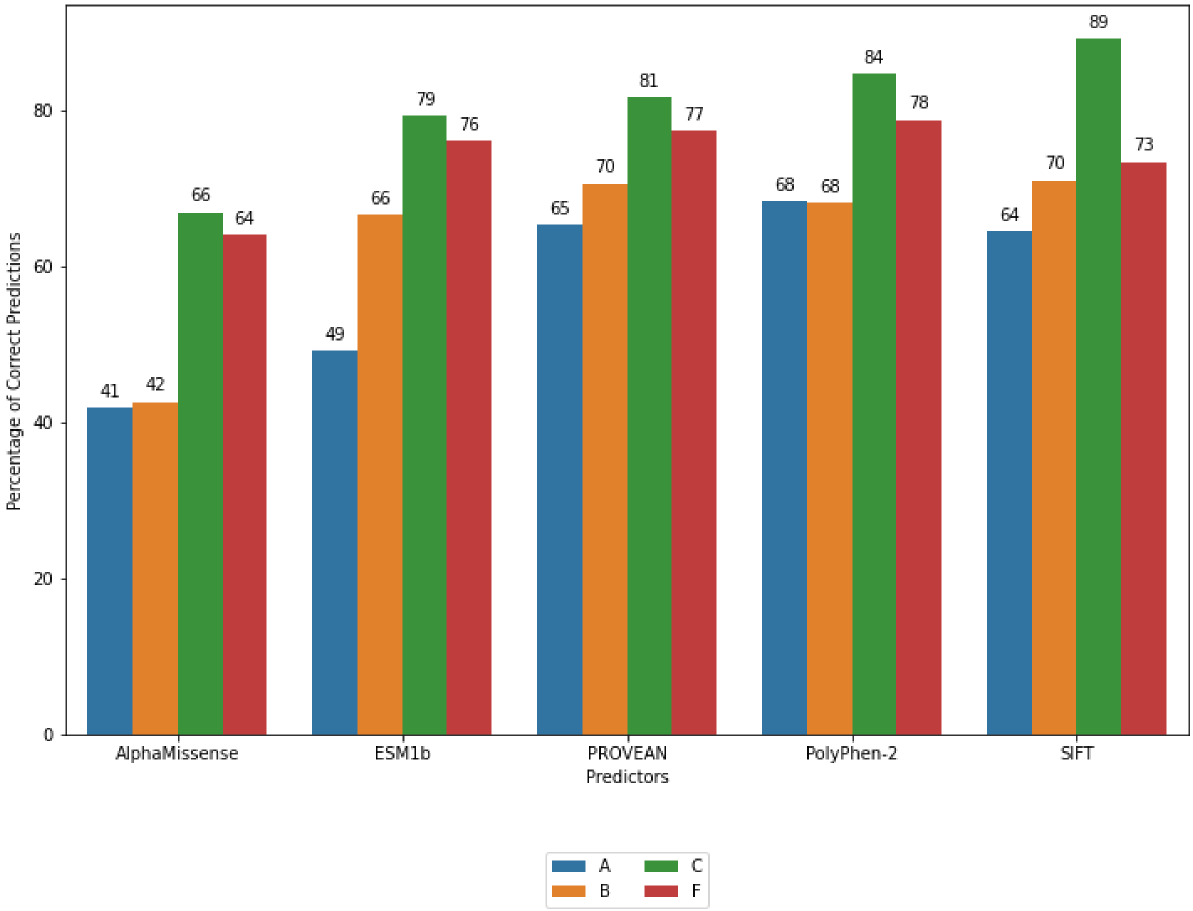


**Figure S12: Bar plot showing the number of correct predictions by predictor and GPCR Classes (MutHTP data set).**

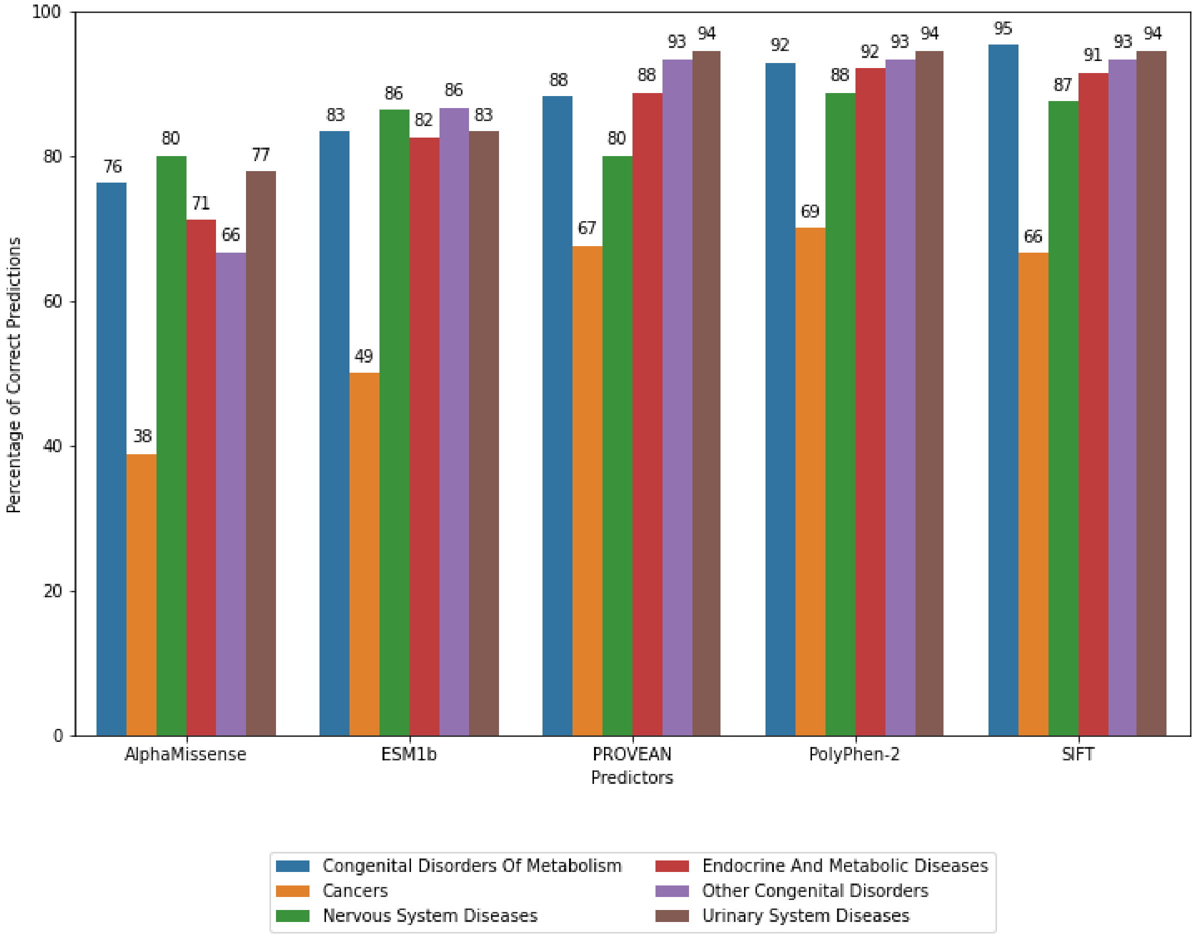


**Figure S13: Bar plot showing the number of correct predictions by predictor and Phenotype, filtered to include only predictors with more than 11 total cases (MutHTP data set).**

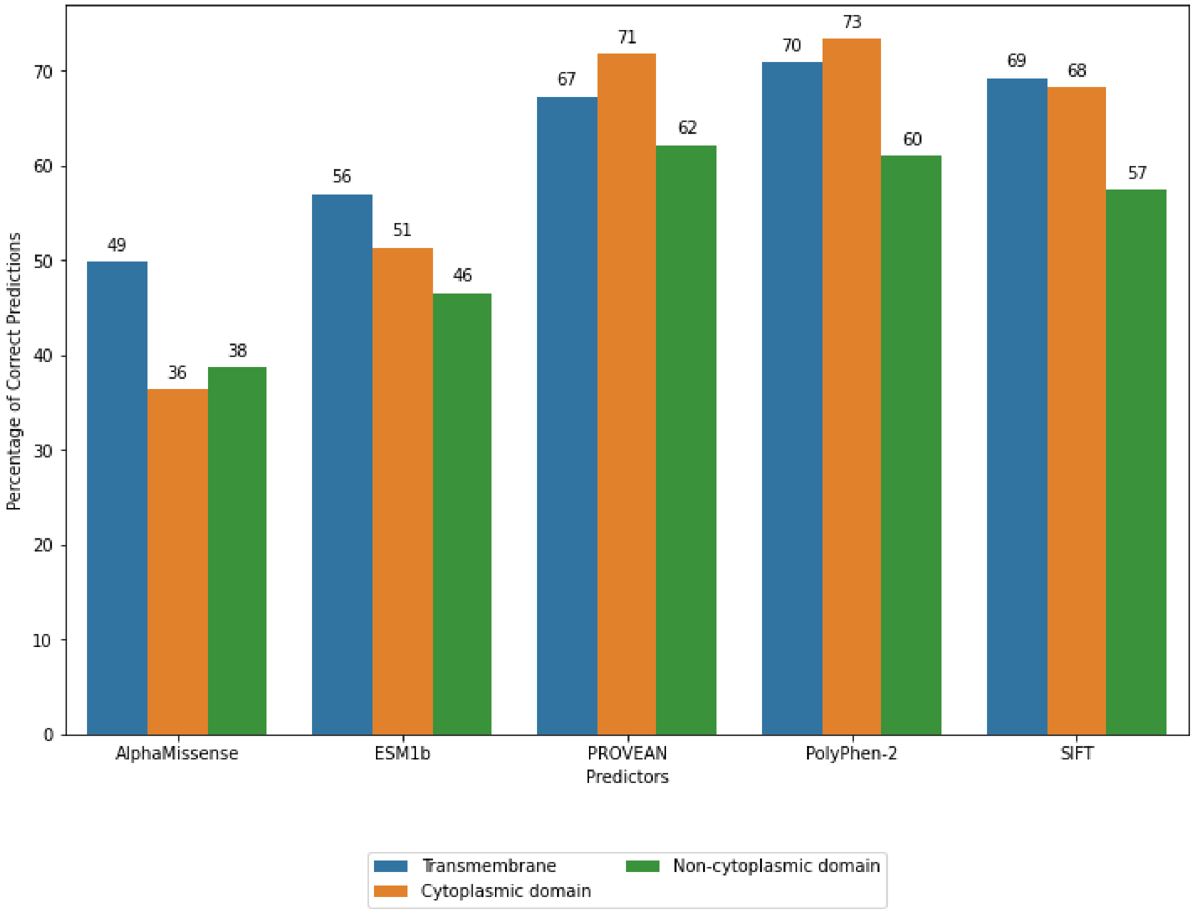


**Figure S14: Bar plot showing the number of correct predictions by predictor and cellular localisation (MutHTP data set).**

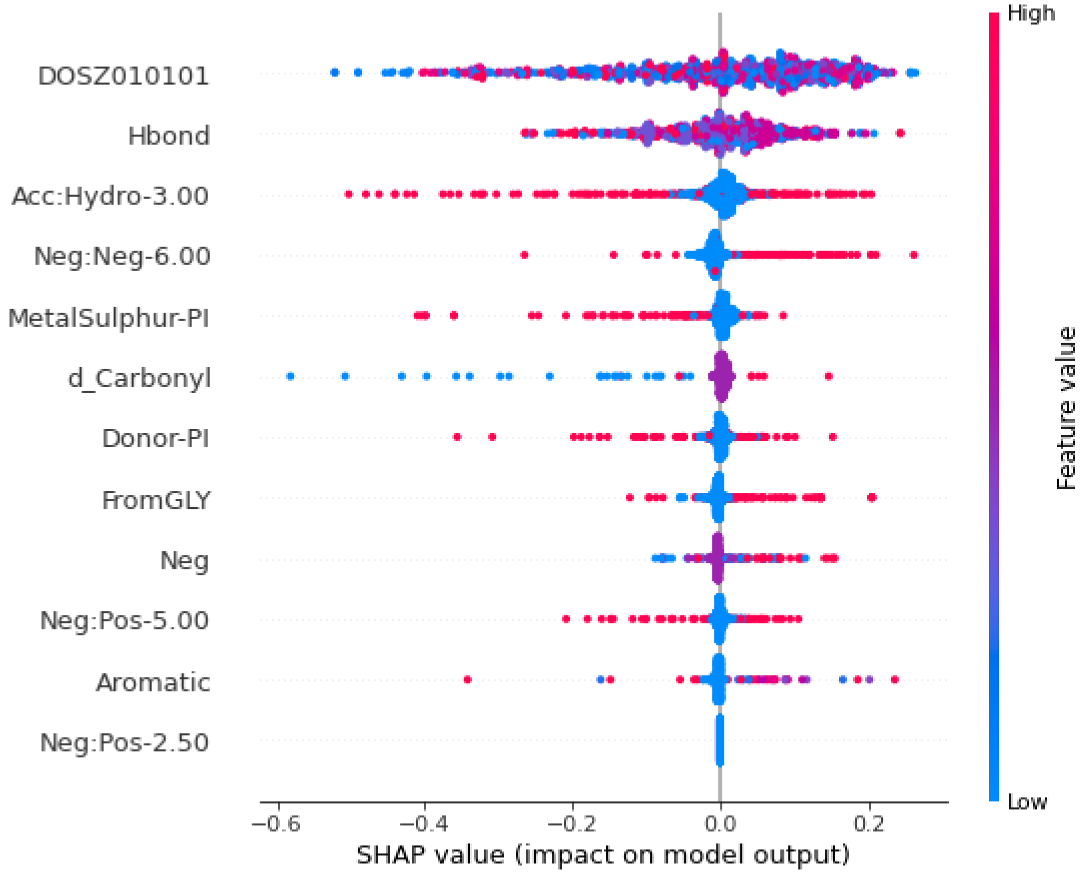


**Figure S15: SHAP feature importance plot for MutHTP dataset. The features that represent higher predictive performance in distinguishing between pathogenic and benign GPCR mutations are represented in descending order of importance on the vertical axis. The points represent the training dataset. High values are represented in red colour, while points with low values are presented in blue. Points to the left influence the prediction to be benign (negative SHAP values), and points, to the right, are pathogenic (positive SHAP values). Different input features affect the output of the respective classification.**

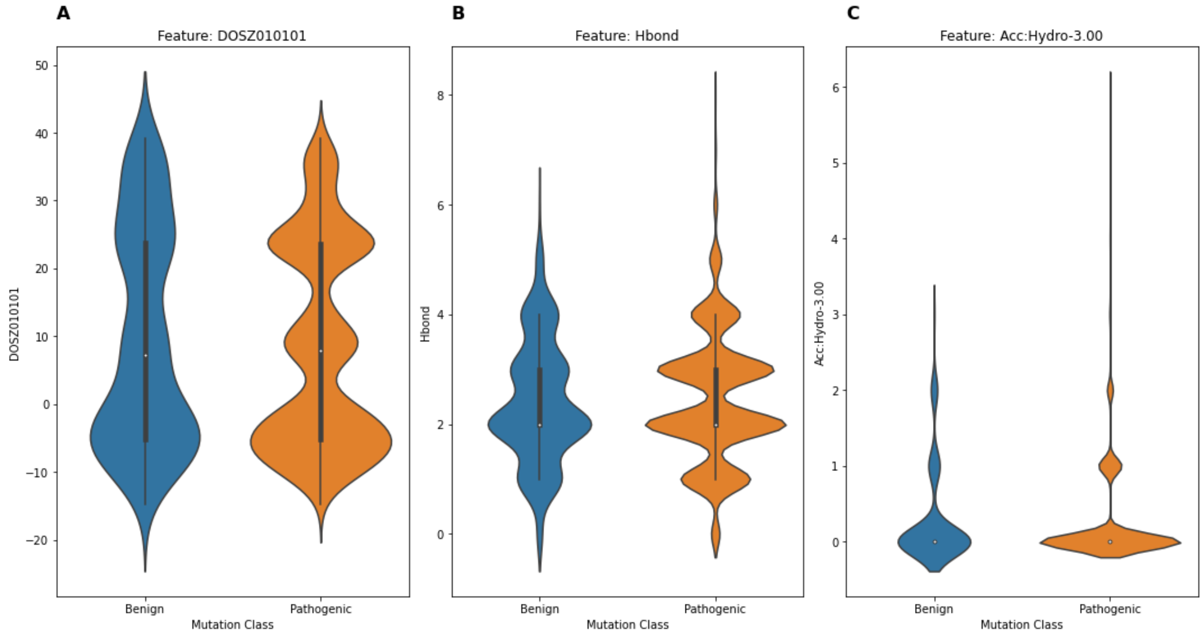


**Figure S16: Violin plot representing the best 3 features when analysing the MutHTP dataset. The violin plot displays the distribution of the 3 features (A: "DOSZ010101", B: "Hbond", and C: "Acc:Hydro-3.00") values for mutation classes. Pathogenic mutations are represented by orange, while benign mutations are represented by blue. The width of the plot at each value indicates the density of data points. Blue represents benign, orange, pathogenic.**

### REFERENCES: SUPPLEMENTARY MATERIAL AND METHODS

Adzhubei, I. A., Schmidt, S., Peshkin, L., Ramensky, V. E., Gerasimova, A., Bork, P., Kondrashov, A. S., & Sunyaev, S. R. (2010). A method and server for predicting damaging missense mutations. *Nat Methods*, *7*(4), 248-249. https://doi.org/10.1038/nmeth0410-248

Boniecki, M., Rotkiewicz, P., Skolnick, J., & Kolinski, A. (2003). Protein fragment reconstruction using various modeling techniques. *J Comput Aided Mol Des*, *17*(11), 725-738. https://doi.org/10.1023/b:jcam.0000017486.83645.a0

Dosztanyi, Z., & Torda, A. E. (2001). Amino acid similarity matrices based on force fields. *Bioinformatics*, *17*(8), 686-699. https://doi.org/10.1093/bioinformatics/17.8.686

Hanahan, D., & Weinberg, R. A. (2011). Hallmarks of cancer: the next generation. *Cell*, *144*(5), 646-674. https://doi.org/10.1016/j.cell.2011.02.013

Jubb, H. C., Higueruelo, A. P., Ochoa-Montano, B., Pitt, W. R., Ascher, D. B., & Blundell, T. L. (2017). Arpeggio: A Web Server for Calculating and Visualising Interatomic Interactions in Protein Structures. *J Mol Biol*, *429*(3), 365-371. https://doi.org/10.1016/j.jmb.2016.12.004

Kawashima, S., Pokarowski, P., Pokarowska, M., Kolinski, A., Katayama, T., & Kanehisa, M. (2008). AAindex: amino acid index database, progress report 2008. *Nucleic Acids Res*, *36*(Database issue), D202-205. https://doi.org/10.1093/nar/gkm998

Luthy, R., McLachlan, A. D., & Eisenberg, D. (1991). Secondary structure-based profiles: use of structure-conserving scoring tables in searching protein sequence databases for structural similarities. *Proteins*, *10*(3), 229-239. https://doi.org/10.1002/prot.340100307

Ng, P. C., & Henikoff, S. (2001). Predicting deleterious amino acid substitutions. *Genome Res*, *11*(5), 863-874. https://doi.org/10.1101/gr.176601

Overington, J., Donnelly, D., Johnson, M. S., Sali, A., & Blundell, T. L. (1992). Environment-specific amino acid substitution tables: tertiary templates and prediction of protein folds. *Protein Sci*, *1*(2), 216-226. https://doi.org/10.1002/pro.5560010203

Pires, D. E., Ascher, D. B., & Blundell, T. L. (2014). mCSM: predicting the effects of mutations in proteins using graph-based signatures. *Bioinformatics*, *30*(3), 335-342. https://doi.org/10.1093/bioinformatics/btt691

Quail, D. F., & Joyce, J. A. (2013). Microenvironmental regulation of tumor progression and metastasis. *Nat Med*, *19*(11), 1423-1437. https://doi.org/10.1038/nm.3394

Skjaerven, L., Yao, X. Q., Scarabelli, G., & Grant, B. J. (2014). Integrating protein structural dynamics and evolutionary analysis with Bio3D. *BMC Bioinformatics*, *15*(1), 399. https://doi.org/10.1186/s12859-014-0399-6

Vogelstein, B., Papadopoulos, N., Velculescu, V. E., Zhou, S., Diaz, L. A., Jr., & Kinzler, K. W. (2013). Cancer genome landscapes. *Science*, *339*(6127), 1546-1558. https://doi.org/10.1126/science.1235122
